# Prediction of small-molecule partitioning into biomolecular condensates from simulation

**DOI:** 10.1101/2025.03.04.641530

**Authors:** Alina Emelianova, Pablo L. Garcia, Daniel Tan, Jerelle A. Joseph

## Abstract

Predicting small-molecule partitioning into biomolecular condensates is key to developing drugs that selectively target aberrant condensates. However, the molecular mechanisms underlying small-molecule partitioning remain largely unknown. Here, we first exploit atomistic molecular dynamics simulations of model condensates to elucidate physicochemical rules governing small-molecule partitioning. We find that while hydrophobicity is a major determinant, solubility becomes a stronger regulator of partitioning in more polar condensates. Additionally, more polar condensates exhibit selectivity toward certain compounds, suggesting that condensate-specific therapeutics can be engineered. Building on these insights, we develop minimal models (MAPPS) for efficient prediction of small-molecule partitioning into biologically relevant condensates. We demonstrate that this approach reproduces atomistic partition coefficients in both model systems and condensates composed of the low complexity domain (LCD) of FUS. Applying MAPPS to various LCD-based condensates shows that protein sequence can exert a selective pressure, thereby influencing small-molecule partitioning. Collectively, our findings reveal that partitioning is driven by both small-molecule–protein affinity and the complex interplay between the compounds and the condensate chemical environment.

## INTRODUCTION

Biomolecular condensates are membrane-less intracellular compartments composed of proteins and nucleic acids and encompass structures such as the nucleolus, stress granules, P-bodies, and centrosomes [1–5]. Many condensates are formed and maintained through phase separation, appearing as dynamic, liquid-like assemblies with distinct chemical environments [6–11]. Such an organization facilitates efficient biochemical reactions and regulatory processes. For example, condensates play essential roles in RNA metabolism, signal transduction, and cellular stress responses [2, 12]. It is, therefore, unsurprising that the dysregulation of biomolecular condensates has been linked to several human diseases. In neurodegenerative disorders, pathological phase separation can lead to the formation of insoluble aggregates. For example, TDP-43 mislocalization and aggregation are hallmarks of amyotrophic lateral sclerosis (ALS), and tau protein aggregates are central to Alzheimer’s disease [13, 14]. In cancer, altered phase separation can drive oncogenesis by enhancing signaling pathways, modulating gene expression, or creating environments favorable to cancer progression [15]. Additionally, many viruses exploit cellular condensates to enhance their replication or shield themselves from immune responses, as seen in certain stress granules or nucleolar condensates [16, 17]. Given these adverse effects, condensates represent promising targets for drug development.

Therapeutic strategies targeting biomolecular condensates are an emerging area of drug discovery, as small molecules can prevent the formation of pathological condensates by disrupting the weak multivalent interactions driving intracellular phase separation [18–21]. The current predominant approach in the field involves a phenotype-based strategy, in which extensive high-throughput screens of small organic molecules are carried out experimentally [22, 23]. Using such screens, certain small molecules have been identified that partition into condensates and modulate their material properties [24–28]. Although potential drug candidates have been identified, the phenotype-based strategies used are inherently agnostic to the mode of action of the drugs. Thus, at present, drug discovery for condensates possesses an inherent blind spot that significantly hampers progress in the field: the molecular mechanisms underlying the preferential partitioning of small molecules into condensates and their consequent effects on condensates remain largely unknown.

The challenge of targeting condensates is further exacerbated by their inherently dynamic nature. Many proteins involved in condensate formation lack well-defined structures, making them unsuitable for traditional structure-based drug design [29–31]. Additionally, variability in condensate chemical environments [19, 32]—i.e., distinct from one condensate to another and determined by their unique compositions—poses an additional challenge in drug discovery. For instance, identifying molecules that target one type of condensate without affecting others requires a precise understanding of the unique interactions within them.

Recent experimental efforts combined with machinelearning-based computational approaches have aimed to uncover the physicochemical factors that govern smallmolecule partitioning into condensates [18]. Thody et al. [33] identified key features of small molecules, such as solubility and hydrophobicity, that favor or disfavor condensate partitioning, which varied over nearly six orders of magnitude. They concluded that the enrichment is primarily driven by the formation of hydrophobic environments within condensates and emphasized the hydrophobicity of a compound as the dominant driver of its partitioning into condensates.

Kilgore et al. [32] probed selective small-molecule partitioning into condensates formed by MED1, NPM1, and HP1a, revealing distinct partitioning patterns across these environments. Using machine learning, they further analyzed how molecular properties such as size, hydrophobicity, flexibility, and number of hydrogen bonds influence partitioning. Their results suggest that the chemical environments of condensates selectively concentrate small molecules based on a complex interplay of physicochemical properties rather than a single dominant feature, including interactions with a specific condensate environment. Additionally, Li et al. [34] showed that variations in the microenvironment heterogeneity of condensates significantly affect the potency of small molecules. Their findings suggest that condensates provide a unique chemical environment, challenging the idea that physicochemical properties of a compound alone dictate partitioning.

Molecular dynamics (MD) simulations offer a compelling approach to understanding the complex relationships between small molecule properties and their selectivity for particular condensates. Crucially, MD simulations enable the capture of the fully disordered nature of condensates while dynamically probing their interactions with small molecules. A recent study by Zhang et al. [35] utilized all-atom MD simulations to explore how non-specific hydrophobic interactions within MED1, HP1*α*, and NPM1 condensates influence the partitioning of small molecules. Their simulations used systems representative of condensates and revealed that the unique compositions and physicochemical properties of the condensates favor certain small molecules over others. While such studies offer valuable insights into the intricate mechanisms of molecular interactions, modeling a phase boundary of a condensate could provide a broader picture of partitioning mechanisms, which is computationally expensive for large systems at the all-atom resolution. In contrast, coarse-grained (CG) models simplify system representation by grouping atoms into interaction sites or “beads”. This reduction in complexity and computational expense enables the screening of a broader range of small molecules, complementing all-atom studies by uncovering larger-scale dynamics.

Here, we employ atomistic and CG molecular dynamics simulations to investigate the factors influencing smallmolecule partitioning into condensates. At an atomistic resolution, we design four model protein condensates [36] with varying physicochemical properties, specifically a decreasing degree of polarity. Using these model systems, we examine the partitioning behavior of 17 organic small molecules spanning a range of physicochemical characteristics, including hydrophobicity, solubility, flexibility, and size (Fig. 1). Together, these ∼65 systems enable a detailed analysis of the interplay between small molecule properties and the chemical environment in determining localization to condensates. Building on insights from the atomistic simulations, we develop a set of minimal CG models to further elucidate the key properties driving small-molecule partitioning. This approach, dubbed MAPPS for MinimAl models for Prediction of small-molecule Partitioning into biomolecular condensateS, enables the efficient exploration of interactions between small molecules and condensates. Together, our work uncovers critical features and empirical relationships that describe interactions between small molecules and condensates: (1) the extent of a compound’s partitioning is driven by the unique characteristics of the condensate environment and is shaped by the physicochemical properties of the compound, (2) the degree of partitioning in more hydrophobic condensates is proportional to the compound’s affinity for the protein chains, which is determined by the specificity of its interactions with protein residues, (3) the capacity of condensate to accommodate the molecules from the dilute phase by effectively excluding water is a strong contributing factor determining the distribution of small molecules across phases.

**FIG. 1:**
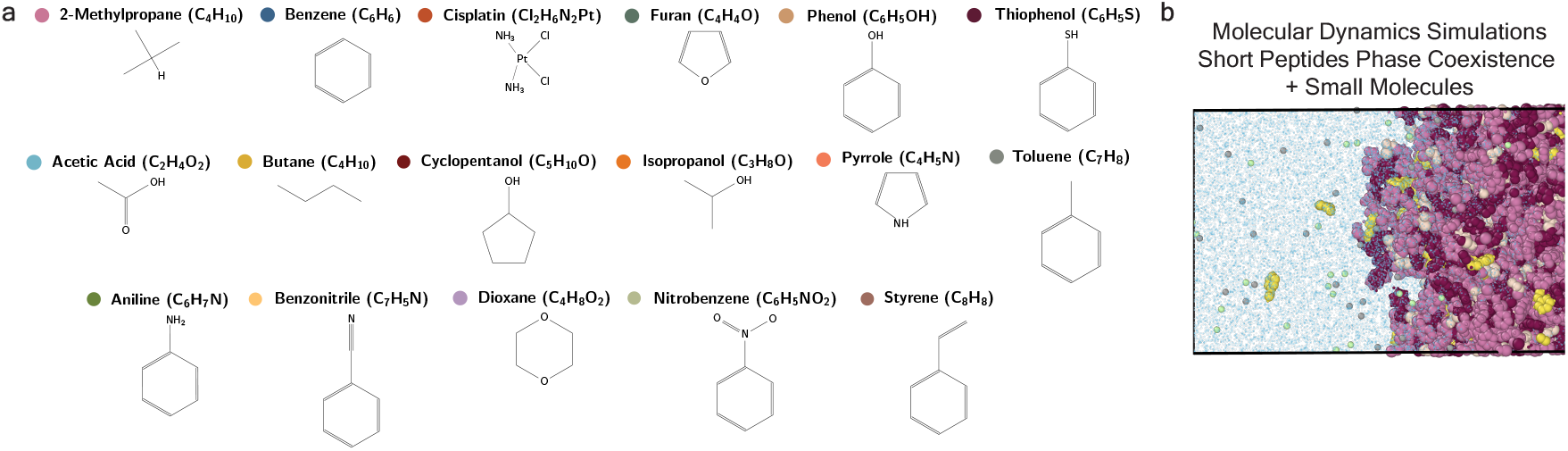
Atomistic molecular dynamics simulations were used to probe the partitioning of organic molecules into model condensates. (a) Chemical structures of the simulated compounds. Colored circles in the titles correspond to the markers’ colors for the data points reported throughout the work. The molecules are ordered alphabetically. (b) Snapshot from the all-atom molecular dynamics simulations (part of the simulation box is shown), illustrating the partitioning of styrene into an FYAFYF model condensate (protein is colored in pink colors, small molecules are in yellow, water is in blue and salt ions are in green and grey).

## RESULTS

### Partitioning is influenced by the physicochemical properties of both small molecules and condensates

To gain atomistic insight into the partitioning of small molecules in various condensates, we simulate ∼65 systems composed of small molecules and model peptide-based condensates (FWAFWF, FYAFYF, YQHQHY, NYANYN). We then calculate the small-molecule partitioning coefficients (*K*) into the protein-rich phase. Fig. 2 shows *K* values versus (a) the compounds’ log *P* values (the partitioning coefficient of a solute between octanol and water, obtained from Xlog *P*3 3.0 [37]), (b) the logarithm of solubility (at 25^°^C based on Ref. [38]), and (c) molecular diameter estimated based on the van der Waals volume using ProteinVolume 1.3 tool [39].

**FIG. 2:**
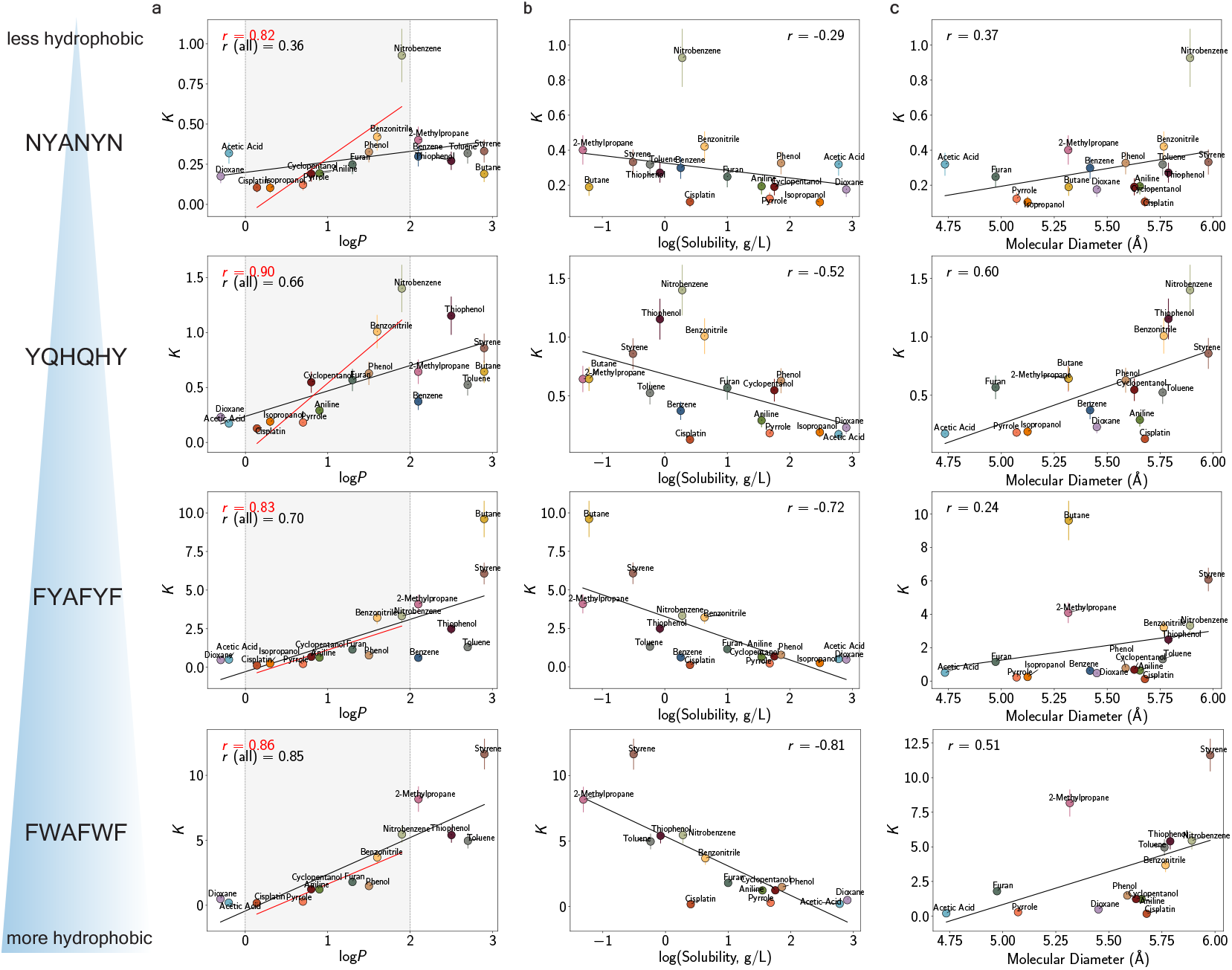
Physicochemical properties of small molecules, as well as chemical environment of condensates, dictate partitioning. Partitioning coefficient of a set of small molecules into model condensates composed of: NYANYN (first row), YQHQHY (second row), FYAFYF (third row), and FWAFWF (fourth row) peptides, based on all-atom simulations at 300 K, plotted against the compounds (a) log *P* values (partition coefficient between octanol and water [37]), (b) solubility (log_10_) at 25^°^C [38], and (c) molecular diameter. The solid black line shows a linear fit with a corresponding Pearson correlation coefficient, and the red solid line shows a linear fit in the region of 0 *<* log *P <* 2. Error bars represent standard errors. The concentration of the small molecules in each system was 100 mM, the protein concentration was 300 mM for FYAFYF, NYANYN, and YQHQHY peptides and 240 mM for FWAFWF peptides.

Atomistic simulations for all peptides reveal distinct partitioning trends: *K* values significantly increase with higher log *P* values, overall ranging from 0.2 to over 12 (Fig. 2a). Hydrophobic molecules, such as styrene and butane, partitioned significantly more into the highly hydrophobic condensates FYAFYF and FWAFWF, with *K* reaching up to

12.5 versus *K ≈* 0.2–0.5 in the polar condensates. This is consistent with previous studies where it was experimentally observed that small molecules preferentially partitioned into condensates with higher hydrophobicity [34].

Another finding was the higher *K* values of dioxane and acetic acid—both with low log *P* values—in all systems, compared to some molecules with higher log *P*, such as isopropanol, pyrrole, and cisplatin. For acetic acid, higher *K* can be attributed to its molecular size: the small size, with an estimated molecular diameter of less than 4.8 Å (Fig. 2c), facilitates easy penetration into the condensate. The higher *K* values of dioxane may be attributed to the more flexible cyclic ether structure compared to the structures of molecules of smaller size and higher log *P* (pyrrole), which are more rigid due to their delocalized π electrons. In contrast, cisplatin, a bulky and rigid metal complex, has limited flexibility due to its platinum center and coordinated ligands, restricting its access to certain regions of the condensate. Although these structural properties have a weaker effect than hydrophobicity and solubility, molecular size and flexibility can still contribute to condensate selectivity, albeit to a lesser extent, consistent with Ref. [32].

While peptide-based model condensates serve as useful systems for understanding the partitioning behaviors, it is important to probe these effects in more biologically relevant condensates. All-atom simulations, while detailed, are computationally expensive and inefficient for exploring a wide variety of systems or capturing the effects of small molecules over long timescales. To overcome these limitations, we leveraged atomistic simulations to develop a minimal coarse-grained approach that focuses on the key factors influencing small-molecule partitioning. This method enables the study of small-molecule partitioning for more biologically relevant condensates, such as those composed of low-complexity domains of proteins.

### Designing minimal models for predicting small-molecule partitioning into biomolecular condensates

The interactions between small molecules and condensates are governed by a complex interplay of forces— including dispersive forces, electrostatics, and hydrogen bonding. Accurately modeling these interactions across a wide parameter space and diverse systems can quickly become computationally prohibitive. To efficiently simulate the partitioning behavior of small molecules and their impact on condensate properties, it is beneficial to use an approach that simplifies the system while retaining essential features. Such an approach should, therefore, effectively describe interactions between small molecules (SM) and macromolecules (e.g., proteins) in condensed systems, while also considering other physical characteristics of condensates, such as solvent-occupied volume.

Several chemically specific CG models have been previously developed to describe protein–protein interactions in phase-separated systems [40–46]. Here, we adopt the Mpipi model [41]—a residue-resolution approach with implicit solvent that has been successfully employed to characterize phase behavior of disordered proteins. The Mpipi potential captures both bonded and non-bonded protein interactions, including long-range electrostatics and shortrange non-bonded contacts. The non-bonded interactions were parameterized using all-atom potentials-of-mean-force (PMF) calculations of amino acid pairs [41, 47] and bioinformatics data [48]. Adopting Mpipi as a foundational model, we develop a minimal CG approach to predict smallmolecule partitioning into condensates (Fig. 3). We hypothesize that this approach will provide an efficient means of estimating the partition coefficients of small molecules into biologically relevant condensates while enabling us to identify the dominant physicochemical properties.

**FIG. 3:**
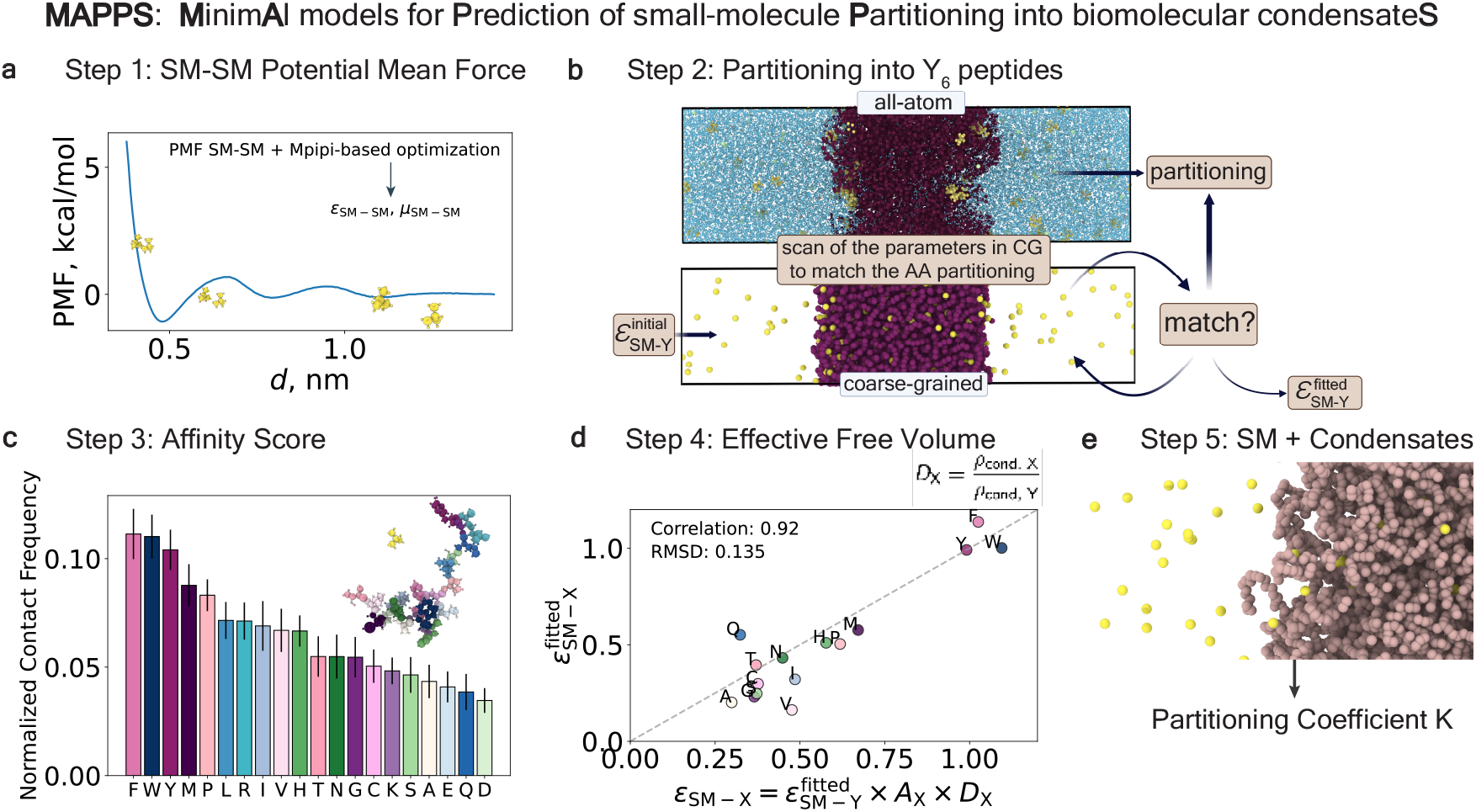
Illustration of the parametrization algorithm used to develop MAPPS. Results for 2-methylpropane are shown as an example. (a) The potential of mean force for a molecule self-interaction was used to determine molecule self-interaction parameters of the Wang-Frenkel potential. (b) A series of slab simulations with Y_6_ peptides at 300 K and salt concentration of 150 mM in coarse-grained resolution was used to optimize the molecule-tyrosine interaction parameter, 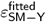, to reproduce partitioning behavior observed in the all-atom simulations of the corresponding system. (c) The distribution of the normalized contacts frequencies between the molecule and a single protein chain, obtained from all-atom simulations at 300 K and salt concentration of 150 mM, was used to derive interaction parameters for the molecule with other residues (*ε*_SM−X_). (d) Validation of the optimization based on the contacts distribution: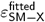 values corresponding to partitioning in all-atom simulations obtained by extending the optimization procedure through all-atom slab simulation in (YXY)_2_ peptides for each residue (X) vs. *ε*_SM−X_ values obtained through Eq. 1. (e) Snapshot from a coarse-grained molecular dynamics simulation illustrating partitioning of the molecules into an LCD condensate, simulated with parameters obtained via steps 1-4.

Since in Mpipi the protein–protein interactions are partially derived from PMF calculations, one approach to incorporating small molecules is by computing PMFs (for SM– SM as well as SM–X pairs, where X represents a protein residue). Here, the compounds in our test set are small enough to be represented as a single interaction site (molecular diameter less than 6 Å) and all have a net zero charge. Thus, PMF calculations (Fig. 3a) can be directly used to determine the parameters representing the relative strength of interactions of the Wang–Frenkel potential [49] which captures short-ranged non-bonded interactions. We initially employed this approach; however, the resulting model failed to accurately reproduce the partitioning observed in our atomistic simulations (data not shown).

There are several reasons why relying solely on SM– SM/SM–X interactions via PMFs is insufficient to capture the observed trends in partitioning. While PMFs can reflect the effective relative interaction strength, they do not effectively capture the effects of a condensate environment, which can affect the available space for the molecules and hence their distribution between phases,–factors important for the partitioning of the compounds. These environmental factors modulate a molecule’s preference between solvent and condensate, making it essential to consider the full context of the protein condensate to accurately describe partitioning behavior.

Additionally, by reducing the complexity of a system by simplifying its representation, the chemical specificity of a compound may be lost due to the exclusion of certain degrees of freedom and the neglect of charge distribution. To enhance the chemical specificity of the model, one approach is to increase the resolution of the SM representation. While this would improve accuracy, it would also result in a higher computational cost. Alternatively, chemical specificity can be captured indirectly by accounting for the key factors that drive small-molecule partitioning into condensates.

Therefore, rather than relying solely on PMFs for parameterization of both SM–SM and SM–X interactions, we design an optimization approach that accounts for both SM characteristics and the physicochemical environment of the condensates. First, as before, we compute PMFs for SM– SM interactions (Fig. 3a), which are used to parameterize the parameters of the Wang–Frenkel potential *ε*_SM−SM_ and *µ*_SM−SM_ (see Eq. 7 in Methods) representing the SM–SM interaction strength in the CG model (see Methods). However, to determine SM–X interactions, we initially design model (YXY)_2_ condensates (where Y represents tyrosine and X represents an amino acid) at atomistic resolution. For a given small molecule, we then characterize the partitioning atomistically and iteratively optimize the CG parameters to reproduce the atomistic observables, effectively determining the interaction terms for SM–X (similar to Fig. 3b). Note that for certain X residues, (YXY)_2_ does not form condensates; for such systems, we use a random forest model (Fig. S7) to derive parameters associated with such residues as part of our parameterization procedure. Strikingly, the resulting CG model is transferable. When we probe SM partitioning at the CG level for condensates (not included in the training set), we observe good agreement with the all-atom behaviors. This finding supports our hypothesis that both small molecule properties and condensate chemical environments are critical for governing partitioning. We should note, however, that the developed models are designed to reproduce partitioning rather than the bulk-phase thermodynamic properties of pure compounds. As a result, the optimization strategy prioritized partitioning behavior over accurately capturing pure-phase properties of the compounds.

A key drawback of the above parameterization approach is its labor-intensive nature. For each new small molecule, the process requires simulating its partitioning into approximately 20 model (YXY)_2_ condensates, simulating the corresponding CG model condensates, and performing optimizations across multiple parameter sets. However, motivated by the success of this optimization approach in capturing partitioning trends, we sought to identify empirical relationships that describe SM–X interactions in condensate environments.

### Deriving an empirical relationship for predicting small molecule–protein interactions in phase-separated systems

Both our optimization approach and original atomistic investigations highlight two key factors that govern small molecule–protein interactions in phase-separated systems: (1) the averaged interactions between small molecules and constituent proteins based on their chemical composition, and (2) interactions between small molecules and the local condensate environment, influenced by factors such as hydrophobicity, polarity, pore size, and water content within the condensates. Combined, these two sets of properties define an effective SM–X interaction between a given small molecule and protein residue X in a phase-separated system. Thus, we define an ansatz that describes this effective interaction, denoted *ε*_SM-X_:

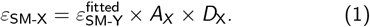

Eq. 1 defines an effective interaction between a given small molecule and a protein residue in a phase-separated system, *ε*_SM-X_, as a function of the effective interaction between the small molecule and tyrosine, 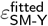. This interaction is scaled by the affinity score for residue X, *A*_X_, and the condensate density ratio for X, *D*_X_—two metrics that we introduce. The first term in Eq. 1, 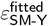, sets a consistent baseline for calibrating interactions involving other amino acids. We define the affinity score of residue X,

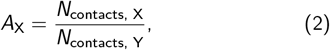

as the likelihood that a given small molecule will bind to residue X in a random peptide chain (Fig. 3c). *A*_X_ is computed by simulating (at atomistic resolution) an ensemble of peptide chains, each 20 amino acids long and composed of one of each amino acid, and calculating the contact frequency between the small molecule and the amino acids, which is then normalized by the contact frequency with a tyrosine residue. We find that computing *A*_X_ for about eight random chains each simulated for 500 ns is sufficient to obtain good statistics (Fig. S8). *A*_X_, therefore, provides a computationally inexpensive (compared to PMF calculations), dynamic, and effective means of estimating the binding strength between a small molecule (SM) and a protein residue (X)—essentially accounting for the chemical specificity of the interaction. Because the diffusion of molecules is influenced by the presence of solvent within the condensate and its ability to move through a condensate network— a factor not captured well in a CG model based solely on pairwise interactions at infinite dilution—Eq. 1 includes an additional factor, *D*_X_, that accounts for the solvent volume fraction in the condensate. Here, we define *D*_X_ as

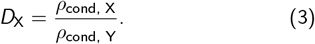

*D*_X_ is calculated by normalizing the density of a model (YXY)_2_ condensate by that of a Y_6_ condensate. Thus, *D*_X_ captures the effective free volume in the condensate when X is present.

Using Eq. 1, we can then determine the effective interaction parameters that describe small molecule–protein interactions in condensates.

### Parameterizing effective interactions between small molecules and proteins in condensates

In this section, we detail the parameterization process using 2-methylpropane as an example (Fig. 3). Other small molecules in this study are parameterized using a similar procedure; the details are provided in the Supporting Information.

The first step is to determine the parameters that define the strength of the self-interactions for a compound. These Wang–Frenkel potential parameters *ε*_SM−SM_ and *µ*_SM−SM_ are calculated based on the SM–SM PMF (described in the Methods section). Fig. 3a illustrates the PMF obtained for a pair of 2-methylpropane molecules (PMFs for the other molecules are shown in Fig. S1).

Next, the effective interaction between the small molecule and tyrosine 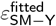 is determined, using optimization through direct coexistence simulations in the slab geometry [44] (Fig. 3b and the Methods section). Here, the initial value for the optimization 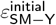 is computed based on the SM–Y PMF. The final value, 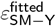 is obtained by optimizing 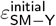 through a series of CG simulations of a Y_6_ condensate, to reproduce the partitioning obtained from all-atom simulations in the corresponding system (Table S4 and Fig. S4c). Rather than performing a PMF calculation to find 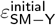, we find that initial values within a range of 0.2 kcal/mol (for less partitioned compounds) to 0.5 kcal/mol (for higher partitioned compounds) work well for all the small molecules tested.

The affinity score *A*_X_ for the SM–X pairs is then computed from Eq. 2 (Fig. 3c) based on generic protein chain simulations (see the Methods section). Finally, the condensate effective free volume parameter *D*_X_ is calculated via CG slab simulations using Eq. 3 (Table S7). These steps yield the final set of effective interaction parameters *ε*_SM−X_, based on Eq. 1.

We verified the applicability of Eq. 1 by omitting the *A*_X_ and *D*_X_ in Eq. 1 and obtaining the parameters for other residues 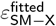 through performing the optimization with (YXY)_2_ peptides, similar to the procedure for obtaining 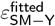. Fig. 3d displays the obtained set 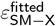 versus _ε SM−X_ obtained by applying Eq. 1. The nearly linear relationship, with a correlation coefficient of 0.92 and rootmean-square deviation (RMSD) of 0.12, suggests that Eq. 1 is a valid approximation to determine the set of parameters for various SM–X pairs based on a single optimization procedure for the SM–Y pair, the affinity score *A*_X_ that captures the chemical specificity of the interactions between a given small molecule and a protein residue, and the condensate density ratio *D*_X_ that implicitly accounts for residue(X)-specific effects on the condensate. The final set of parameters for all small molecules (i.e., MAPPS) are provided in Table S8.

### Small-molecule partitioning observed atomistically is captured via coarse-grained simulation

#### Peptide-based condensates

The MAPPS approach was initially validated by simulating the partitioning of selected small molecules into model condensates. For five peptide-based condensates (FYAFYF, FWAFWF, YQHQHY, NYANYN, and NFWAFS), we simulated a subset of small molecules that exhibited divergent partitioning in all-atom simulations: cisplatin, phenol, 2-methylpropane, styrene, and nitrobenzene. The comparison of the partitioning coefficients obtained from CG simulations via MAPPS with the corresponding values from the atomistic simulations is presented in Fig. 4a. The CG simulations capture the trends observed in the atomistic simulations for each model condensate, showing increased partitioning with increasing compound hydrophobicity. Overall, we observe better agreement with the all-atom simulations for more hydrophobic condensates: FWAFWF and FYAFYF. For FWAFWF, there is almost quantitative agreement reached for all compounds. Based on these results, we additionally verified the performance of the models for a range of the concentrations of the compounds using the FWAFWF system (Fig. S10).

**FIG. 4:**
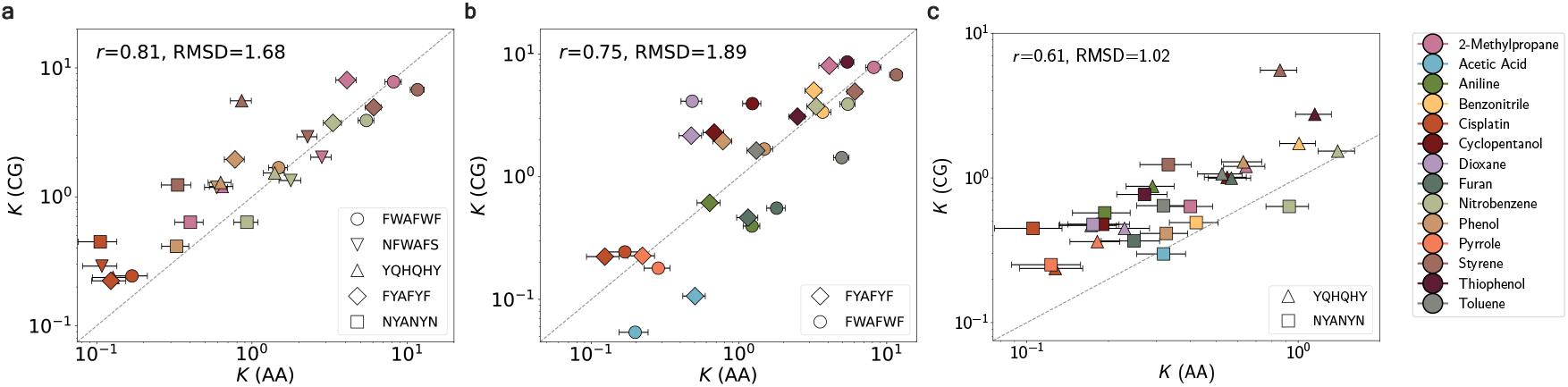
Validation of transferability of MAPPS on systems of short peptides. Coarse-grained (CG) and all-atom (AA) simulations of a condensate with small molecules were conducted at 300 K and 150 mM salt concentration, with the protein concentration of 300 mM for FYAFYF, NYANYN, and YQHQHY peptides and of 240 mM for FWAFWF peptides, and the small molecules concentration of 100 mM. (a) Partitioning coefficients of a subset of small molecules (cisplatin, phenol, 2-methylpropane, styrene, and nitrobenzene) in the CG simulation versus AA simulations, for a set of peptides of various degrees of polarity FWAFWF, NFWAFS, YQHQHY, FYAFYF, NYANYN; (b) partitioning of a larger set of small molecules in the CG simulation versus AA simulation for FYAFYF and FWAFWF peptides and (c) YQHQHY and NYANYN peptides. The Pearson correlation coefficient *r* and the root-mean-square deviation (RMSD) values are displayed. The dashed line represents the equity line.

For the larger set of compounds (displayed in the legend of Fig. 4), we simulate their partitioning in FYAFYF, FWAFWF, YQHQHY, and NYANYN condensates, and compare the results against the values obtained in the corresponding all-atom simulations (Fig. 4b,c). The simulations capture the increasing magnitude of the *K* values with increasing peptides hydrophobicity, ranging from the maximum of 1.2 for NYANYN, 2.8 for YQHQHY, and reaching approximately 8 in FYAFYF and FWAFWF. Notably, the agreement between the results from the two resolutions was stronger for compounds partitioning into the more hydrophobic condensates, FYAFYF and FWAFWF (Fig. 4b). Fig. S9 additionally shows the CG results plotted similarly to Fig. 2 for *K* versus log *P*, solubility, and the molecular diameter (size). The CG simulations correctly predict diminished partitioning for thiophenol and toluene compared to 2-methylpropane and nitrobenzene in FYAFYF. The CG simulations also exhibit quantitative predictions for almost all the systems with benzonitrile, nitrobenzene, and phenol. Additionally, the preference for compounds such as nitrobenzene (lower hydrophobicity) over 2-methylpropane (higher hydrophobicity) was captured for YQHQHY (Fig. 4c).

In contrast to the all-atom simulations, the molecular size emerges as a more significant factor governing partitioning in the CG simulations (with consistently higher correlation coefficients, Fig. S9). Additionally, the CG simulations systematically overestimate partitioning for most of the compounds in the polar condensates, in particular for high log *P >* 2 compounds such as styrene. The discrepancy comes from the fact that highly hydrophobic compounds with low solubility, such as styrene, exhibit significantly lower partitioning into polar condensate environments, which may not be captured as effectively via MAPPS. We hypothesize that the predictions for such systems may be improved by additional terms in Eq. 1 that account for molecule solubility. For all condensates, a significant deviation from the atomistic predictions was observed for dioxane, cyclopentanol, furan, and acetic acid. Of note, these are all oxygen-containing molecules; thus, points to a need to incorporate additional adjustments in the procedure to account for the specific chemistry of such interactions.

In summary, MAPPS captures the trends in the partitioning values for hydrophobic environments but yields a systematic overestimation of partitioning into more polar condensates. The reduced accuracy of the approach in polar environments suggests a deficiency in accounting for small molecule–solvent interactions. This highlights the need for a refined parameterization that incorporates additional interaction types to more faithfully account for the effect of the solvent.

#### FUS LCD condensate

In addition to the peptide-based condensates, we also test our MAPPS approach against atomistic simulations for condensates made up of low-complexity domains (LCDs). Here, we report our results for small-molecule partitioning into a FUS LCD condensate. To simulate the all-atom (AA) condensate, we first equilibrate a FUS condensate at the CG level using Mpipi [41] and use a backmapping procedure [50] to generate the corresponding AA configuration (Fig. 5a). Due to the high computational costs, the all-atom simulations with FUS include only a small set of compounds— cisplatin, 2-methylpropane, pyrrole, and thiophenol—which we also simulated using the CG approach. Specifically, the compounds chosen for testing give good agreement between CG and all-atom predictions for the peptide-based condensates.

**FIG. 5:**
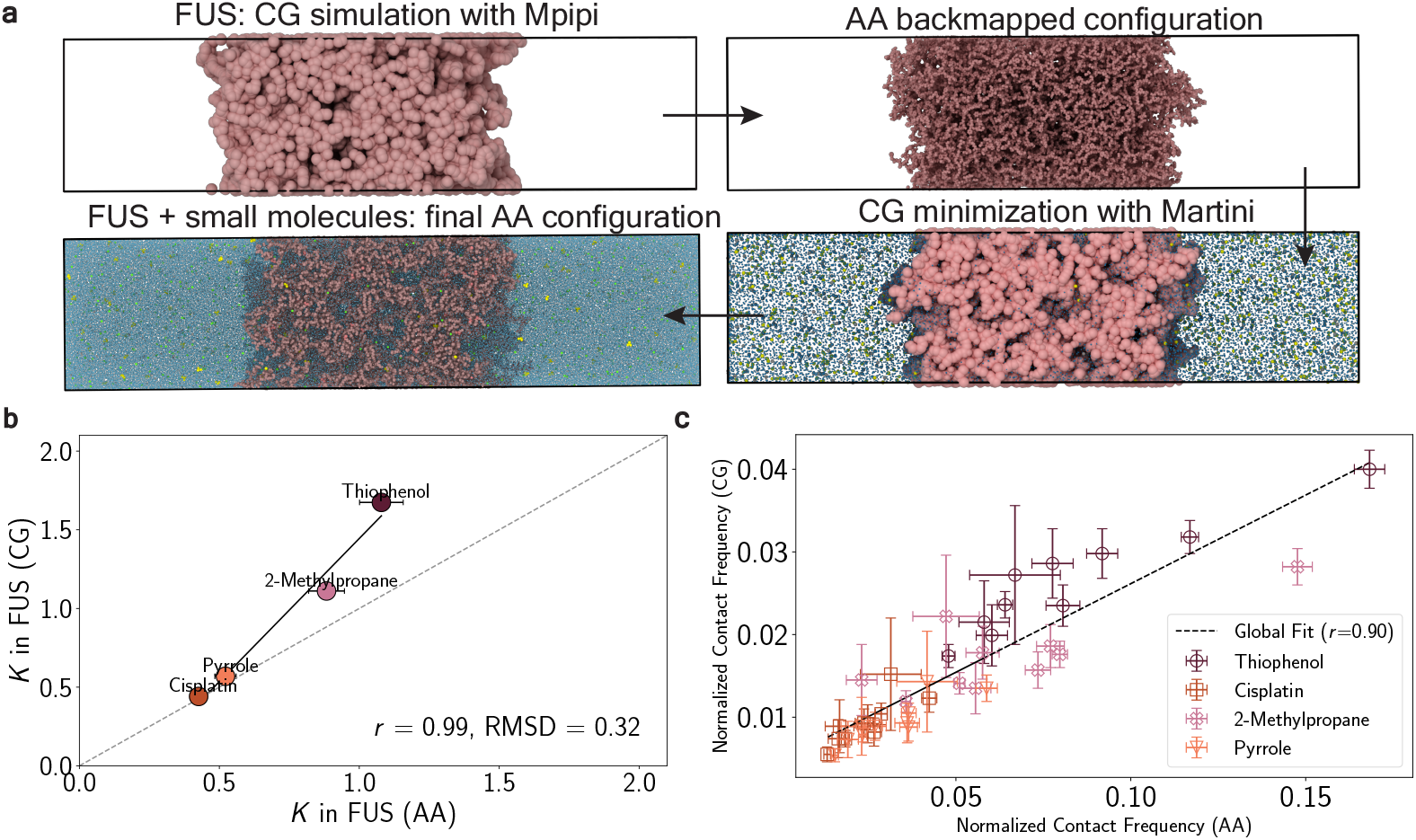
Simulations of FUS LCD condensate with small molecules in all-atom (AA) and coarse-grained (CG) resolutions. Simulations were conducted at 300 K and 150 mM salt concentration. 64 chains of FUS LCD were simulated, at a protein concentration of 12.5 mM and small molecules concentration of 30 mM. (a) Backmapping procedure to obtain the AA configuration of the condensed FUS LCD chains. (b) Partitioning coefficients obtained in CG simulation versus AA simulations. The error bars for the values from the CG simulations are within the markers. (c) Normalized contact frequency between a molecule and a single residue of FUS-LCD in the CG simulation versus AA simulation (each marker corresponds to a single residue). The errors are computed as standard errors for the frequency of contacts across the analyzed time frames.

Across the two resolutions, we obtain a good agreement in partitioning coefficients for the compounds tested (Fig. 5b; *r* = 0.99); although the CG predictions were systematically higher. The quantitative discrepancy may primarily stem from the longer time scales needed to capture the partitioning in the all-atom simulation, where diffusion is hindered by the presence of explicit solvent and ions. Additionally, since the FUS LCD has a large fraction of polar residues (58%), the consistently higher *K* values observed in the CG simulations may result from the systematic overestimation of the partitioning into more polar environments, as was shown for the peptide-based systems.

We also examine whether the interaction landscapes of the compounds in the condensates are consistent at the two length scales. Specifically, we measure the contact probability of each compound with the residues in FUS LCD in the phase-separated systems (Fig. 5c). Here, we observe a strong linear correlation (*r* = 0.90) between the frequency of residue contacts at the AA and CG resolutions. This result strongly suggests that the CG simulations can capture key interaction patterns found in the higher-resolution AA system. Figure S12 also shows the distribution of these values per residue for each of the compounds, illustrating high correlation for each compound.

### LCD-based condensates exhibit varying partitioning of small molecules

Using CG simulations, we investigate small-molecule partitioning into condensates composed of low-complexity domains (LCD) of proteins. We focus on FUS, hnRNPA1 (here, A1), EWSR1, and TIA1 LCDs that have been shown to be important for condensate formation in vitro and incells [51–55]. The small molecule-to-protein molar concentration (in the range of 0.2–0.7) is chosen to be consistent with experimental assays [56, 57]. Fig. 6a illustrates the residue composition of each LCD. Fig. 6b shows the distribution of the partitioning coefficients for each LCD obtained for a set of molecules at 300 K and 150 mM salt. Additionally, Fig. 6c–f depict the *K* values as a function of the compounds log *P* value, for each LCD system.

**FIG. 6:**
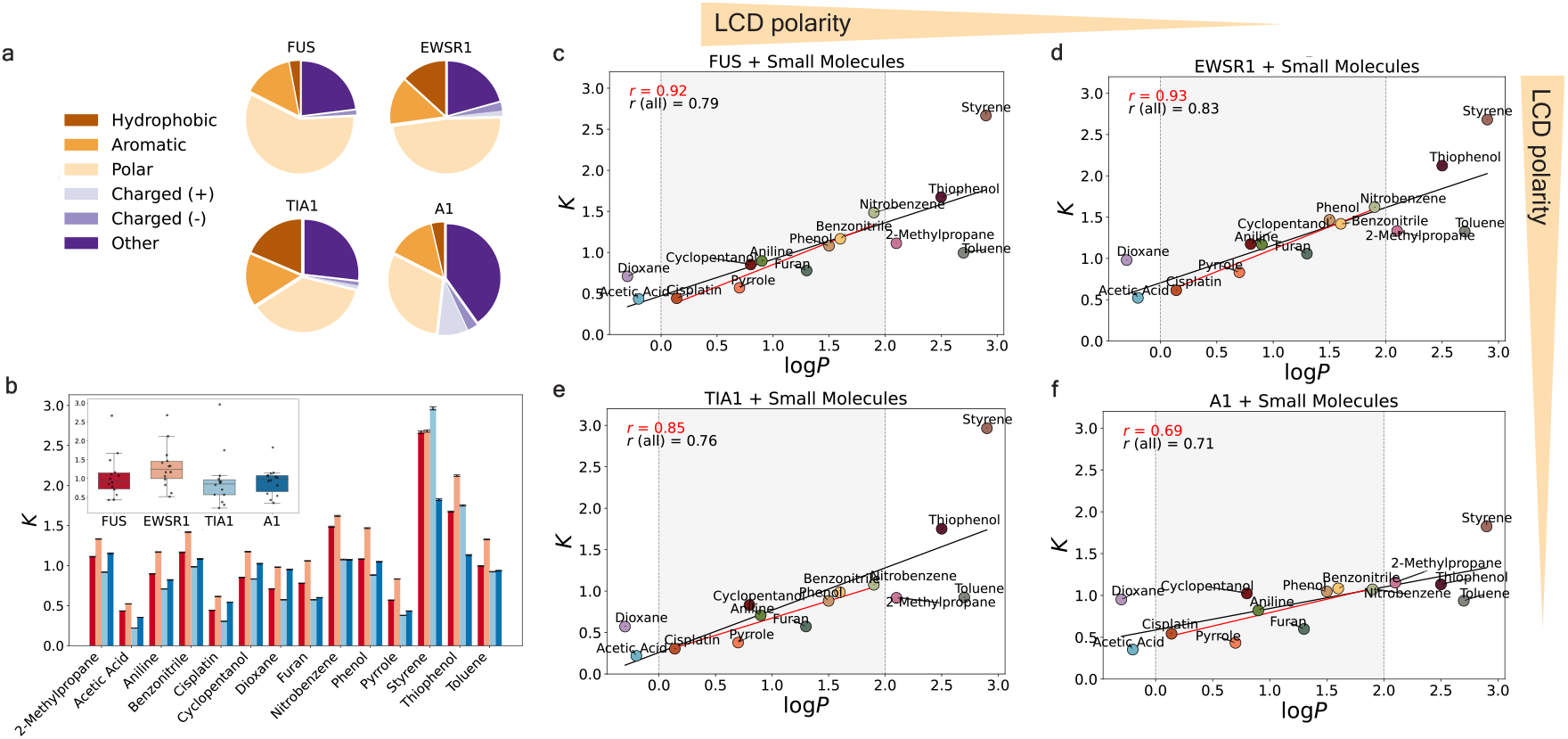
Coarse-grained simulations of small molecules partitioning into condensates of low-complexity domains of proteins. The following low-complexity domains were simulated: FUS, EWSR1, TIA1, and A1 of 12 mM, 7 mM, 20 mM, 16 mM concentrations, respectively. The concentration of the small molecules for each system was 30 mM. Simulations were conducted at 300 K and 150 mM salt concentration. (a) Compositions of the residues for the simulated LCDs. The following residues classification is used: “hydrophobic” – A, I, L, M, V; “aromatic” – F, W, Y; “polar” – N, Q, S, T; “charged (+)” – R, K, H; “charged (-)” – D, E; “other” – C, G, P. (b) Partitioning coefficients *K* for FUS, EWSR1, TIA1, and A1 LCDs per compound. Error bars represent standard errors. The inset plot is the box-and-whisker plot of partitioning coefficients *K* for each protein across all tested molecules. The box represents the interquartile range (IQR), with the lower and upper edges corresponding to the first (Q1) and third (Q3) quartiles, respectively. The horizontal line inside the box indicates the median *K* value. (c-f) Partitioning coefficients plotted against the compounds log *P* values. The yellow arrow represents the decreasing percentage of the polar residues among the LCDs. Displayed values *r* is the Pearson correlation coefficient.

The partitioning coefficients show the greatest variability for hydrophobic compounds, such as styrene, which ranged from *K* = 1.75 in A1 to *K* = 3.0 in TIA1, where styrene achieved its highest partitioning. Thiophenol also displays significant variation, with *K* values ranging from 1.1 in A1 to 2.1 in EWSR1, highlighting the heightened sensitivity of hydrophobic compounds to differences in condensate environments. Nitrobenzene shows a strong preference for FUS and EWSR1, as indicated by its higher *K* values exceeding 1.5. Notably, only in A1 did dioxane exhibit a higher *K* value than furan, suggesting a specific affinity of dioxane for residues in A1 condensates.

To analyze the differences in the overall distribution of the partitioning values among the proteins, we conduct the Kruskal–Wallis test [58]. This non-parametric test is chosen as it does not assume normality and is suitable for comparing multiple independent groups. Following the Kruskal–Wallis test, Dunn’s post-hoc test with Bonferroni correction [59] is performed to assess pairwise differences between the proteins.

The Kruskal–Wallis test results (test statistic *H* = 7.60, with *p* = 0.055) suggest that while some differences may exist in *K* among the proteins, the overall statistical evidence is weak, as the *p*-value is slightly above the conventional significance threshold of 0.05. However, post-hoc pairwise comparisons using Dunn’s test reveals a potentially significant difference between EWSR1 and TIA1 (*p* = 0.046).

A closer examination of partitioning trends indicates that in the EWSR1 system most compounds favor the condensed phase (*K >* 1), with only pyrrole and cisplatin showing a preference for the dilute phase. In contrast, in the other protein systems—FUS, A1, and particularly TIA1—lower partitioning coefficients are observed overall. Notably, only nitrobenzene, styrene, and thiophenol demonstrate a strong preference for the condensed phase in these systems. TIA1, in particular, shows systematically lower partitioning, with a median *K* below 1. This is largely due to the higher density of TIA1 compared to other systems (Fig. S11), which generally leads to more complex diffusion of the compounds. However, when high-affinity interactions are present, they override transport limitations, resulting in higher *K* values compared to less dense condensates, as seen in the case of styrene partitioning in TIA1. Hence, despite the generally lower partitioning in TIA1, compounds with high log *P* values exhibit the highest *K* values in this system, significantly shifting its median partitioning. This pattern arises from the high proportion of hydrophobic and aromatic residues in TIA1, which strongly enhance interactions with highly hydrophobic compounds.

More broadly, our findings demonstrate that partitioning is not solely dictated by chemical specificity but is also shaped by condensate composition and physical properties, particularly density. Thus, accurately comparing different proteins requires recognizing the dynamic balance between diffusion constraints and high-affinity interactions. Hence, factors such as crowders used to form the condensed phase and other environmental factors must be accounted for by experimentalists evaluating molecular potency, since ignoring these considerations may lead to misinterpreting partitioning trends within condensates.

## DISCUSSION

In recent years, biomolecular condensates have been positioned as therapeutic targets, which has sparked intense interest in elucidating the factors that drive small-molecule partitioning into condensates. In this work, we leverage molecular dynamics simulations to interrogate which factors dictate localization of small molecules into condensates. At atomistic resolution, we systematically examine partitioning behaviors across ∼65 systems comprising a diverse set of small molecules and peptide-based condensates. Our findings reveal that for small molecules with low hydrophobicity (log *P <* 2), partitioning into condensates scales approximately linearly with log *P*, particularly in condensates enriched in hydrophobic residues. However, for more hydrophobic compounds (log *P >* 2), this linear relationship weakens, and solubility becomes the dominant factor. We find that this effect is particularly pronounced in condensates enriched in polar residues. Our findings suggest that even when protein condensates are, on average, more hydrophobic than their surrounding environment, the presence of polar residues overrides the expected dependence of hydrophobicity on molecular partitioning, leading to a preferential selection of molecules with lower log *P* values. This highlights the critical role of the condensate chemical environment in dictating molecular partitioning within condensates. Overall, our atomistic simulations are consistent with recent studies [32, 33] and point to a complex interplay between small-molecule characteristics and condensate chemical environment.

Levering these insights, we optimize a set of minimal coarse-grained (CG) models (MAPPS) to efficiently predict small-molecule partitioning into biologically relevant condensates. Our approach, built upon a residue-resolution CG model of proteins [41], captures the chemical specificity of small molecule–protein interaction via an affinity score and accounts for the effective free volume for accommodating small molecules via a derived empirical relation. We show that MAPPS demonstrates good transferability and predictability for different systems, including in predicting small-molecule partitioning into LCD-based condensates, validated by all-atom simulations of the FUS LCD system. MAPPS effectively captures partitioning ratios among various compounds for condensates enriched in hydrophobic residues, and accurately reflects increased partitioning in more hydrophobic condensates. MAPPS, therefore, offers a fast and versatile approach for estimating the partitioning of organic compounds into biologically relevant protein condensates.

As examples, we demonstrate that MAPPS enables efficient screening of small-molecule interactions with condensates of FUS, hnRNPA1, EWSR1, and TIA1 LCDs, revealing key differences in partitioning behavior across these proteins. We find that in EWSR1 condensates, small molecules show strong preference for the condensed phase, whereas partitioning is generally lower in TIA1 condensates. However, highly hydrophobic molecules deviate from this trend, displaying increased partitioning in TIA1, likely due to selective interactions driven by its high proportion of hydrophobic and aromatic residues. The variation in partitioning across different proteins underscores the influence of sequence composition and chemical specificity of small-molecule interactions. Interestingly, despite being the most hydrophobic LCD, TIA1 does not necessarily promote the strongest partitioning. Our simulations indicate that TIA1 condensates exhibit higher densities under similar conditions, resulting in smaller pore sizes and restricted diffusion. This result suggests that beyond chemical affinity, physical properties such as condensate density impose additional constraints on partitioning.

Despite the success of MAPPS in capturing trends in partitioning, we find that as the fraction of polar residues grows, the approach overestimates partitioning relative to all-atom results. Since the LCDs probed here contain a sizable fraction of polar residues, we expect the predictions to preserve the trends in partitioning while representing an upper bound. MAPPS also shows significant deviations for highly-polar small molecules, as well as oxygen-containing molecules such as ethers. These discrepancies highlight opportunities to better account for effects such as solubility and self-interactions of molecules in future refinements.

Together, our modeling approach enables the rapid estimation of partitioning trends in biologically relevant condensates, demonstrating quantitative accuracy for highly hydrophobic compounds in hydrophobic condensates. By following the MAPPS procedure, new compounds of interest can be efficiently parameterized for predictive simulations. In the future, we aim to expand MAPPS to a comprehensive library of parameterized compounds, including practical pharmaceuticals, providing a readily accessible resource for screening interactions with LCDs.

More broadly, our findings demonstrate that chemical properties of compounds do not solely dictate smallmolecule partitioning but are also shaped by condensate physicochemical properties. This finding underscores the importance of considering both molecular interactions and condensate properties when evaluating partitioning behavior. Consequently, experimentalists must account for factors such as crowding agents and condensate density when assessing the selectivity of small molecules for condensates.

## METHODS

### Simulations models

#### Mpipi model for protein–protein interactions

The coarse-grained (CG) models for the small molecules are built upon Mpipi [41], a residue-resolution sequence-specific CG model, which was previously shown to effectively predict phase separation of intrinsically disordered proteins. In Mpipi, each amino acid is represented by a single bead, with its corresponding physical properties such as mass, molecular diameter (*σ*), charge (*q*), and an energy scale that reflects the relative frequency of planar π–π contacts (*ε*). The total interaction energy is a sum of bonded interactions within a chain *E*_bond_, electrostatic contributions *E*_elec_, and short-ranged non-bonded interactions between beads *E*_pair_:

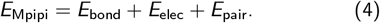

Bonds in the Mpipi model are represented via harmonic springs:

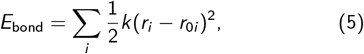

where *k* is the force constant and *r*_0*i*_ is the equilibrium distance. *k* is set to 0.192 kcal/mol/Å^2^ and *r*_0*i*_ to 3.81 Å. The electrostatic contribution is modeled via a Coulomb interaction term with Debye–Hückel screening:

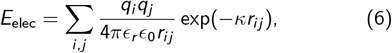

where *ϵ*_*r*_ = 80 is the relative dielectric constant of water, *ϵ*_0_ is the electric constant, and *κ*^−1^ = 0.795 nm is the Debye screening length, corresponding to a monovalent salt concentration of 150 mM. The uncharged non-bonded interactions are modeled via the Wang–Frenkel potential:

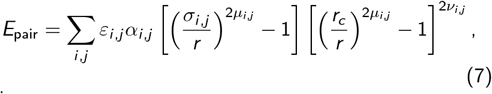

where

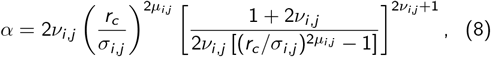

and *σ*_*i,j*_, *ε*_*i,j*_ and *µ*_*i,j*_ are parameters specified for each pair of interacting beads, and *r*_*c*_ is the cutoff radius. For most amino acids, in the original Mpipi implementation, *µ*_*i,j*_ = 1 and *ν*_*i,j*_ = 1. *r*_*c*_ is 3*σ*_*i,j*_ for each pair. All parameters for the Mpipi model can be found in Ref. [41].

#### Coarse-grained parameters for small molecules

The atomistic potential-of-mean-force (PMF) profiles are used to determine the relative strength of self-interactions of the small molecules (*ε*_SM-SM_), described by the Wang– Frenkel potential (Eq. 7). The area of the first well in PMF profiles for a pair of molecules is numerically evaluated and compared to the PMF of tyrosine–tyrosine interactions. This ratio is then used to assess the relative strength of the interactions between the molecules. The parameters *ε*_SM-SM_ and *µ*_SM-SM_ (with fixed *ν*_SM-SM_ = 1 according to the value for most amino acids in Mpipi) are then varied so that (1) the ratio *ε*_SM−SM_ to *ε*_Y_ (tyrosine self-interaction from the Mpipi model) deviated by no more than 10% from the ratio of a minimum of the PMF profile of the molecule– molecule interaction to the minimum of the PMF profile of the tyrosine–tyrosine interaction, (2) the ratio of the well areas under the curve of the PMF profile to such area of the Wang–Frenkel potential corresponding to the SM–SM and tyrosine–tyrosine pairs deviated by no more than 10%. The obtained parameters *ε*_SM-SM_, *µ*_SM-SM_ are assigned to the small molecule represented as a single interaction site (a bead). The bead size *σ*_SM−SM_ is calculated based on the molecule’s estimated van der Waals radius. Each bead is assigned a neutral charge. The *σ*_SM−X_, which represents the small-molecule–X (where X is an amino acid) distance when the potential is equal to zero, is obtained based on an arithmetic rule, using the values of the Mpipi model for the amino acids *σ*_residue_.

### Test systems

We study a set of small molecules to assess the impact of hydrophobicity and solubility on the partitioning: 2-methylpropane, acetic acid, aniline, benzene, benzonitrile, butane, cisplatin, cyclopentanol, dioxane, furan, isopropanol, nitrobenzene, phenol, pyrrole, styrene, thiophenol, toluene. These molecules are selected as they are comparable in size and small enough to represent as a single bead (estimated molecular diameter less than 6 Å) and have zero net charge. The transferability of the models is validated by calculating their partitioning into a variety of model condensates: FWAFWF, FYAFYF, NYANYN, NFWAFS, and YQHQHY. The number of protein chains, small molecules and their molar concentration in each system are given in Table S3 for the all-atom simulations and Table S6 for the CG simulations.

We also examine partitioning of small molecules into condensates formed by the low-complexity domain (LCD) of the proteins FUS, EWSR1, TIA1, and A1 of 12 mM, 7 mM, 20 mM, 16 mM concentrations, respectively. The concentration of each of the small molecules is 30 mM. The choice of the concentrations is based on the recent experimental works on small-molecule partitioning, where the molar concentration of small molecules was several times higher than the protein [56, 57], and to ensure a consistent number density of protein residues in a simulation of approximately 1.2 beads/nm^3^. All systems are modeled using direct coexistence simulations in the slab geometry.

### Potential-of-mean-force calculations

PMF calculations are performed with the GROMACS software [60]. The Amber03ws force field with the TIP4P water model is used to model amino acids and water–ion interactions [61]. The initial structures of the small molecules are optimized with the Restrained Electrostatic Potential (RESP) fitting approach [62] at the HF/6-31G(d,p) level of theory using Gaussian software [63]. The charges obtained are combined with the GAFF2 force field [64] parameters for bonded and non-bonded interaction parameters of a molecule, generated via AmberTools [65]. As GAFF2 does not incorporate parameters for platinum, the AMBER force field parameters for cisplatin are taken from Ref. [66].

Each pair of molecules is placed in a cubic box of size 40 Å × 40 Å× 40 Å and solvated with water and ions at 150 mM NaCl concentration. The distance between the centers of mass of the two structures is used as the umbrella sampling coordinate, with a distance of 14 Å taken as the reference state. At least 50 umbrella windows are used for each pair, with additional windows added for certain systems that require extra sampling. Each window is initially energy-minimized using the steepest-descent algorithm. An umbrella sampling simulation of 10 ns is performed for each window, with the harmonic force constant varying from 2000 to 6000 kJ/mol/nm^2^. The potential mean force profile is then calculated using the weighted histogram analysis method [67].

### Frequency of contacts

Atomistic simulations are used to estimate the normalized number of contacts between a small molecule and each protein residue. To do this, we simulate a set of 7 to 8 single chains composed of the 20 amino acids randomly arranged. Each chain and molecule are placed in a cubic box and solvated with water and ions. The Amber03ws force field with TIP4P water is used. The system is energy-minimized, and a simulation in the *NV T* ensemble is performed for at least 500 ns. The number of contacts between the nonheavy atoms of a residue and the molecule within 6 Å is calculated from the trajectories. First, contact values are binarized (1 if present, 0 otherwise), then summed to obtain the total number of frames with contacts. This sum is divided by the total number of frames, yielding a normalized contact frequency between 0 and 1. For each system, we perform three independent runs, placing the molecule in a random position. The mean and standard error of the mean are calculated across the chains and the three sets of runs for each chain.

In the all-atom simulation of the FUS LCD condensate, the contacts are computed for trajectories from 300 ns to 600 ns of the simulation time, collected every 10 ns. The values are additionally normalized by the number of occurrences of the residue in the protein sequence and the total chain length, and the error is estimated as the standard error across the values.

In the CG simulations of the FUS LCD condensate, the number of contacts is averaged across the trajectories collected every 1 ns, calculated within a pair-wise cutoff distance for each residue–small molecule pair (*r*_cut_ = 1.5 × *σ*_SM−X_). A contact is defined as any occurrence of the center of mass of a molecule within the cut-off distance from a residue’s center of mass. The normalized contact frequency values are calculated using a method similar to that applied for the all-atom simulation data.

### Simulation of phase coexistence and partitioning

#### All-atom simulations of peptide-based condensates

All-atom simulations of phase coexistence in a solution with small molecules are performed using GROMACS software [60]. Small molecules are represented as in the PMF calculations. For protein structures, the Amber03ws force field is used with the TIP4P water model [61]. The initial configurations of the peptides (capped with the NME and ACE groups) are generated using PyMOL [68].

For the model peptides, 100 to 150 chains are simulated. For each system, the peptides are first placed in a cubic box, minimized with the steepest descent algorithm, and the system was relaxed in *NPT* ensemble at 1–50 bar and 300 K for 50–100 ns with a time step of 1–2 ps, to achieve the protein density of approximately 1 g/cm^3^. The temperature is kept constant using a stochastic velocity rescaling algorithm and the pressure is kept constant with a Parrinello–Rahman barostat. Bonds involving hydrogen atoms are constrained to their equilibrium lengths using the LINCS algorithm. Long-range electrostatic interactions are modeled using the particle-mesh Ewald method with a grid spacing of 0.1 nm. Dispersion interactions and short-range repulsion are described by a Lennard–Jones potential with a cutoff at 0.9 nm.

The isotropically compressed configuration is then expanded into a slab, with one of the axes of the box elongated. The slab is solvated with water and ions, and the systems are energy minimized and further equilibrated in the *NV T* ensemble—yielding a peptide-rich (condensed) phase and a peptide-poor (dilute) phase in coexistence. For each system, small molecules are added to the box (dilute phase region). The box is further solvated with water and ions at 150 mM concentration. The obtained configurations are energy minimized and simulated in the *NV T* ensemble, with equilibration for 500 ns and production runs of at least 500 ns to obtain an equilibrated cumulative average of the partitioned molecules (evolution of the values with time is shown in Fig. S4).

For estimation of the partitioning coefficients, we analyze trajectories from the production runs, where data points are collected at 10 ns intervals. The number density profiles across the long axis of the box (divided into 50 slices, which was verified to be optimal) are averaged over the collected trajectories (Fig. S4). The location of the interface (and hence the condensate) is determined using the DBSCAN (Density-Based Spatial Clustering of Applications with Noise) algorithm [69]. Partitioning coefficient *K* is estimated as the ratio of the number of molecules (per bin on average across the trajectory) within the condensate to the number of molecules in the dilute phase:

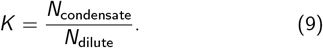

The error associated with partitioning is estimated by propagating the uncertainty from the averaged number density profiles (see Supporting Information for the details on the error analysis).

#### Coarse-grained simulations

All coarse-grained simulations are conducted using LAMMPS (version 23 Jun 2022) [70–73]. The protein interactions are modeled with the Mpipi force field. Initially, the protein copies are compressed in the *NPT* ensemble using a time step of 10 fs and coupled to a Langevin thermostat with a relaxation time of 10 ps. The simulation box was then expanded to a slab configuration, maintaining a volume corresponding to the concentration in the analogous all-atom simulation, achieving a total protein density of approximately 300 kg/m^3^. Each simulation is subsequently run in the *NV T* ensemble for 200 ns to obtain pre-equilibrated condensate configurations. Small molecules are randomly added to the slab configuration of the protein (*N*_total_ = 350 for peptide-based systems and *N*_total_ = 95–150 for LCDs). An *NV T* simulation is then performed for 200 ns of equilibration and 500 ns production, over which the average properties and the standard errors are computed.

The phase boundary of the condensate is estimated based on the slope of the protein density profile. Partitioning is then determined by analyzing the density profile of the small molecules relative to the condensate region. We then calculate the total number of molecules within the “condensate boundary”. As in the all-atom simulations, the partitioning coefficient *K* is calculated as the ratio of the number of molecules (per bin on average across the trajectory) within the condensate boundary to the number of molecules in the dilute phase.

#### All-atom simulations of FUS LCD

The initial structure for all-atom simulations of the phase-separated FUS-LCD system in a slab configuration is mapped from a CG slab simulation with the Mpipi model, which is initially run for 1-microsecond with 64 FUS-LCD chains. To convert the equilibrium CG condensate into fully atomistic systems, we employ a multi-step procedure aimed at accurately reconstructing the atomistic details while minimizing steric clashes [50]. The first step involves reconstructing each chain from the final CG structure in all-atom detail using the Pulchra [74] algorithm, a tool for restoring atomic-level connectivity and geometry from CG models. Pulchra operates by interpreting the CG structure’s backbone and side-chain information, generating atomistic models with proper stereochemistry.

Next, we address the steric clashes that inevitably arise in the reconstructed atomistic structure due to differences in resolution between the CG and all-atom models and the high density of the condensate. To do this, we map the configuration to the Martini2 model [75] using the Martinize script [76]. This step is performed to reduce atomic overlaps while still preserving the explicit representation of backbone and side-chain atoms. In addition, we use standard Martini water and ions to represent the environment.

The third phase of the backmapping process utilizes the “backward” program [77], which is designed specifically for mapping the CG Martini configuration back onto the full atomistic resolution. In this step, we exclude the water molecules from the Martini configuration. The result is a refined all-atom configuration. Finally, we filled the simulation box with water molecules and ions at 150 mM NaCl concentration. The all-atom direct coexistence simulations in the slab geometry are then carried out for 600 ns, using the Amber03ws force field with TIP4P water.

### Performance of MAPPS

The entire MAPPS procedure is described by Eq. 1 and in Fig. 3. Step 1 involves a single PMF calculation for a given small molecule (∼50 GPU hours). Step 3 is the computation of the set affinity scores *A*_X_ pertaining to the small molecule. This can be estimated from about 24 independent (8 chains and 3 random molecule placements) single-chain simulations (total of ∼100 GPU hours). Step 4, which involves the condensate density ratio, does not need to be computed per small molecule. Instead, we provide these values in Table S7 for all X residues. The only expensive part of this method is Step 2 (Fig. 3b). On average, it takes about 7 microseconds of aggregate simulation time: ∼1 microsecond (or 120 GPU hours) for the atomistic *K* and ∼6 microseconds (or 2,500 CPU hours ≈125 GPU hours) in CG for the optimization to get a good estimate of 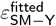. Combining with 1–4, step 5 can then be run for a biologically relevant condensate to estimate *K* in a given system which takes ∼1 GPU hour. With our approach, the entire procedure takes ∼400 GPU hours and allows rapid screening of multiple systems through the CG simulations, while a 1-microsecond all-atom simulation of LCDs used to estimate partitioning in one system requires more than 1000 GPU hours (starting from an equilibrium condensate configuration obtained through a backmapping procedure).

## CODE AND DATA AVAILABILITY

The source code for MAPPS and input scripts used for simulations can be found at the Joseph Group GitHub repository: https://github.com/josephresearch/MAPPS.

## SUPPORTING INFORMATION

Amino-acid sequences of LCDs studied, potential-of-mean-force profiles for small molecule pairs, MAPPS parameters, additional all-atom and coarse-grained simulation results (PDF).

## ACKNOWLEDGMENTS

We thank Nathaniel Hess and Virginia Jiang for their insightful discussions and feedback. We also thank the members of the Joseph Group for their feedback during the development of this approach. This research was partially supported by an Innovation Grant from the Princeton University School of Engineering. J.A.J. also acknowledges research support from the Chan Zuckerberg Initiative DAF (an advised fund of Silicon Valley Community Foundation; grant 2023-332391) and the National Institute of General Medical Sciences of the National Institutes of Health under Award Number R35GM155259. All simulations in this work were performed using the Princeton Research Computing resources at Princeton University, which is a consortium of groups led by the Princeton Institute for Computational Science and Engineering (PICSciE) and the Office of Information Technology.

