## Supplementary File for "Prediction of small-molecule partitioning into biomolecular condensates from simulation"

### Supporting Information for: Prediction of small-molecule partitioning into biomolecular condensates from simulation

[1]*Department of Chemical and Biological Engineering, Princeton University, Princeton, NJ 08544, USA*
[2]*Omenn–Darling Bioengineering Institute, Princeton University, Princeton, NJ 08544, USA*
(Dated: March 4, 2025)

#### CONTENTS

S1. Amino-acid sequences of LCDs studied S1

S2. Potential-of-mean-force profiles for small-molecule pairs S3

S3. Calculation of partitioning S3
A. All-atom simulations S3
B. Coarse-grained simulations S9

S4. MAPPS parameters for small molecules S11

S5. Peptides + small molecules: additional results S17

S6. LCDs + small molecules: additional results S18

References S19

#### S1. AMINO-ACID SEQUENCES OF LCDS STUDIED

Table S1. 3- and 1-letter codes and indexes for the amino acids

| 3-letter code | 1-letter code | Index |
|---|---|---|
| MET | M | 1 |
| GLY | G | 2 |
| LYS | K | 3 |
| THR | T | 4 |
| ARG | R | 5 |
| ALA | A | 6 |
| ASP | D | 7 |
| GLU | E | 8 |
| TYR | Y | 9 |
| VAL | V | 10 |
| LEU | L | 11 |
| GLN | Q | 12 |
| TRP | W | 13 |
| PHE | F | 14 |
| SER | S | 15 |
| HIS | H | 16 |
| ASN | N | 17 |
| PRO | P | 18 |
| CYS | C | 19 |
| ILE | I | 20 |

| | |
|---|---|
| **FUS** | MASND YTQQA TQSYG AYPTQ PGQGY SQQSS QPYGQ QSYSG YSQST DTSGY GQSSY SSYGQ SQNTG YGTQS<br>TPQGY GSTGG YGSSQ SSQSS YGQQS SYPGY GQQPA PSSTS GSYGS SSQSS SYGQP QSGSY SQQPS YGGQQ<br>QSYGQ QQSYN PPQGY GQQNQ YNSSS |
| **A1** | GSMAS ASSSQ RGRSG SGNFG GGRGG GFGGN DNFGR GGNFS GRGGF GGSRG GGGYG GSGDG YNGFG NDGSN<br>FGGGG SYNDF GNYNN QSSNF GPMKG GNFGG RSSGG SGGGG QYFAK PRNQG GYGGS SSSSS YGSGR RF |
| **TIA1** | MINPV QQQNQ IGYPQ PYGQW GQWYG NAQQI GQYMP NGWQV PAYGM YGQAW NQQGF NQTQS SAPWM GPNYG<br>VQPPQ GQNGS MLPNQ PSGYR VAGYE TN |
| **EWSR1** | MASTD YSTYS QAAAQ QGYSA YTAQP TQGYA QTTQA YGQQS YGTYG QPTDV SYTQA QTTAT YGQTA YATSY<br>GQPPT GYTTP TAPQA YSQPV QGYGT GAYDT TTATV TTTQA SYAAQ SAYGT QPAYP AYGQQ PAATA PTRPQ<br>DGNKP TETSQ PQSST GGYNQ PSLGY GQSNY SYPQV PGSYP MQPVT APPSY PPTSY SSTQP TSYDQ SSYSQ<br>QNTYG QPSSY GQQSS YGQQS SYGQQ PPTSY PPQTG SYSQA PSQYS QQSSS YGQQS SFRQD HPSSM GVYGQ |

#### S2. POTENTIAL-OF-MEAN-FORCE PROFILES FOR SMALL-MOLECULE PAIRS

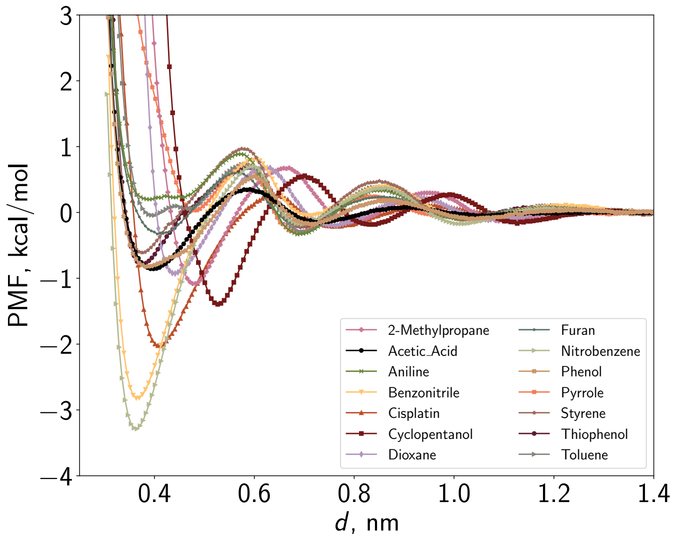


Figure S1. **Potential of mean force (PMF) for molecule–molecule pairs as a function of distance between their center of mass.** The profiles are computed via umbrella sampling simulations at 300 K with GROMACS software. Each pair of molecules is placed in a cubic box of size 40 Å × 40 Å × 40 Å and solvated with water and ions at 150 mM NaCl concentration. The distance between the centers of mass of the two structures is used as the umbrella sampling coordinate, with a distance of about 14 Å taken as the reference state.

#### S3. CALCULATION OF PARTITIONING

##### A. All-atom simulations

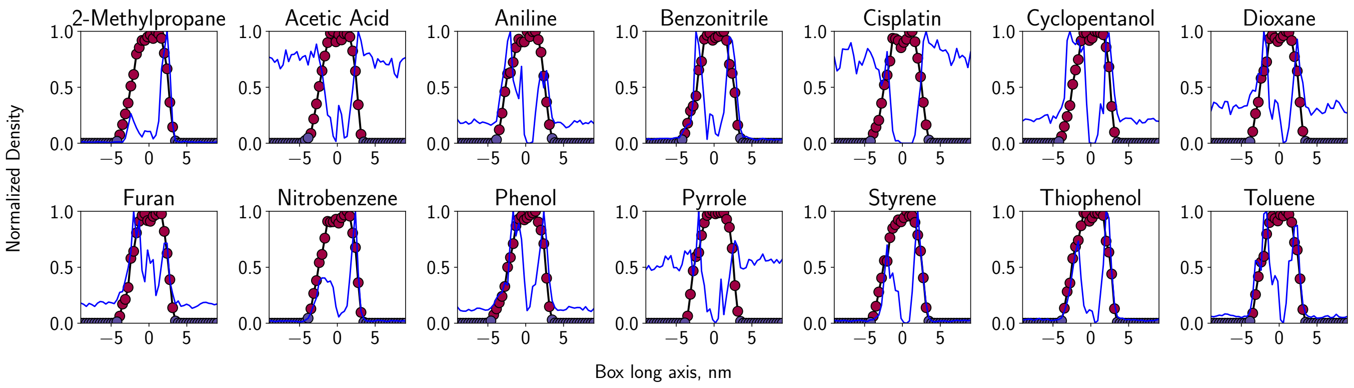


Figure S2. **Density distributions of protein (markers) and small molecules (solid blue line) for the $Y_6$ peptides + small molecules system obtained from the all-atom simulations at 300 K**. Red markers indicate the phase boundary region, within which the small molecule density was used to calculate partitioning. The corresponding partitioning coefficients are provided in Table S4 and are utilized for the parameterization of the compounds.

Table S3. **System parameters for all-atom slab simulations.** Simulations of phase coexistence in a solution with small molecules (denoted here as SM) are performed using GROMACS software [1]. The Amber03ws force field is used with the TIP4P water model [2]. A consistent setup is applied across systems with different small molecules.

| Protein | Box size [Å$^3$] | $N$ SM | $N$ protein chains | $C$ SM [mM] |
|---|---|---|---|---|
| YMYYMY | 226.0 × 55.7 × 55.7 | 50 | 100 | 119 |
| YGYYGY | 284.6 × 49.6 × 49.6 | 50 | 100 | 118 |
| YTYYTY | 234.5 × 54.6 × 54.6 | 50 | 100 | 119 |
| YAYYAY | 204.5 × 51.1 × 51.1 | 50 | 100 | 155 |
| YYYYYY | 184.0 × 55.2 × 55.2 | 50 | 100 | 150 |
| YVYYVY | 254.5 × 52.4 × 52.4 | 50 | 100 | 118 |
| YLYYLY | 224.4 × 52.6 × 52.7 | 50 | 100 | 133 |
| YQYYQY | 227.3 × 55.5 × 55.5 | 50 | 100 | 118 |
| YWYYWY | 193.4 × 56.7 × 56.7 | 50 | 100 | 133 |
| YFYYFY | 208.2 × 54.6 × 54.6 | 50 | 100 | 133 |
| YSYYSY | 238.3 × 54.2 × 54.2 | 50 | 100 | 118 |
| YHYYHY | 233.3 × 54.8 × 54.8 | 50 | 100 | 118 |
| YNYYNY | 201.9 × 55.5 × 55.5 | 50 | 100 | 133 |
| YPYYPY | 230.8 × 55.1 × 55.1 | 50 | 100 | 118 |
| YCYYCY | 252.6 × 52.6 × 52.6 | 50 | 100 | 118 |
| YIYYIY | 255.7 × 52.3 × 52.3 | 50 | 100 | 118 |
| FWAFWF | 259.2 × 52.0 × 52.0 | 50 | 100 | 118 |
| FYAFYF | 250.0 × 58.2 × 58.2 | 50 | 150 | 100 |
| NYANYN | 248.8 × 58.3 × 58.3 | 50 | 150 | 100 |
| NFWAFS | 274.0 × 50.6 × 50.6 | 50 | 100 | 118 |
| YQHQHY | 248.8 × 58.3 × 58.3 | 50 | 150 | 100 |
| FUS | 126.3 × 532.2 × 126.3 | 150 | 64 | 30 |

Table S4. **Partitioning coefficients $K$ for the molecules in a system with $Y_6$ condensate in atomistic simulations.**

| Small Molecule | $K$ |
|---|---|
| 2-Methylpropane | 17.8565 ± 1.7403 |
| Acetic Acid | 0.5984 ± 0.0962 |
| Aniline | 1.7782 ± 0.2341 |
| Benzonitrile | 7.6872 ± 0.9578 |
| Cisplatin | 0.4100 ± 0.0315 |
| Cyclopentanol | 1.9814 ± 0.2459 |
| Dioxane | 1.0451 ± 0.1434 |
| Furan | 2.0926 ± 0.2634 |
| Nitrobenzene | 11.0295 ± 1.1908 |
| Phenol | 3.1884 ± 0.4295 |
| Pyrrole | 0.4864 ± 0.0829 |
| Styrene | 9.2377 ± 1.0471 |
| Thiophenol | 7.8040 ± 0.9622 |
| Toluene | 5.2443 ± 0.6128 |

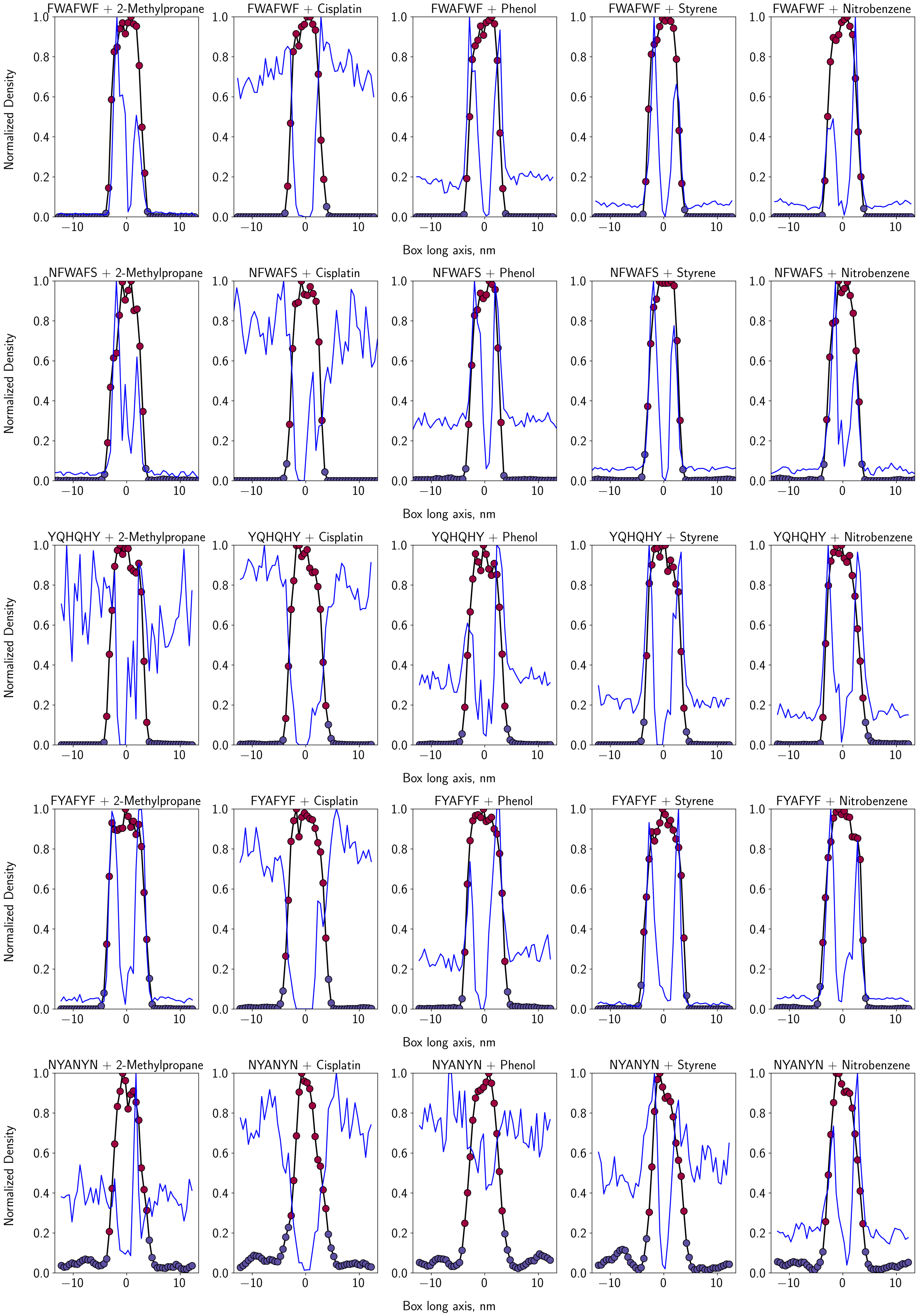


Figure S3. **Averaged density distributions of the protein (markers) and small molecules (solid blue line) for various peptides + small molecule systems obtained from the all-atom simulations at 300 K.** Red markers indicate the phase boundary region, within which the small molecule distribution is counted as partitioning.

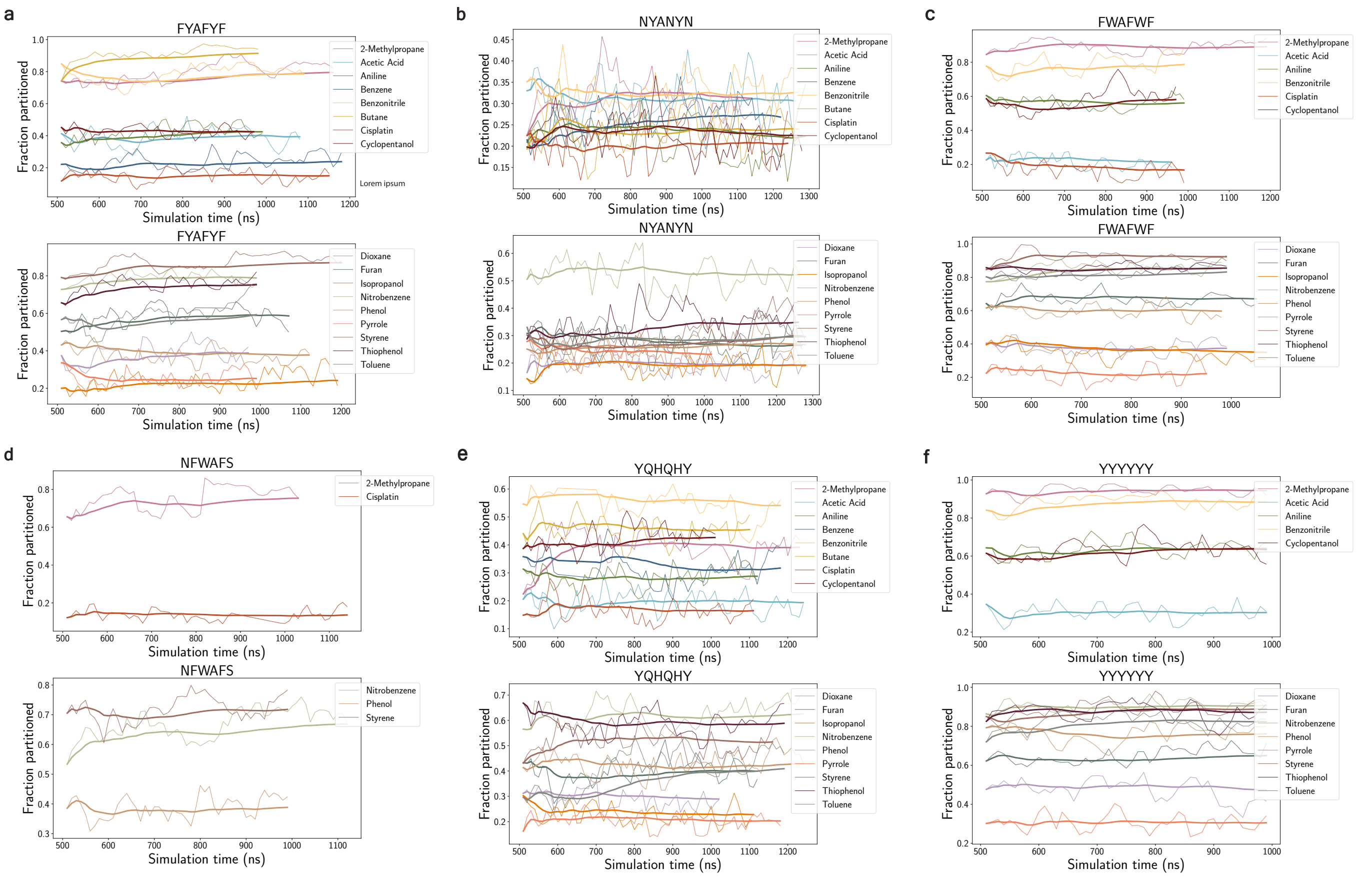


Figure S4. **Fraction partitioned (ratio of the number of small molecules within the condensate $N_{\mathrm{condensate}}$ to the total number of small molecules in the system $N_{\mathrm{total}}$) for the peptide-based systems in the all-atom simulations as a function of the production run time.** The time intervals depicted are used to estimate the average $K$ values. The values are computed every 10 ns (thin lines). The thick lines show the cumulative average.

**Error analysis:** The error of the density distribution for each bin is computed as the standard deviation across the time intervals of 10 ns of a production run. Given a system where the total number of molecules $N_{\text{total}}$ is computed by summing number densities across the number densities computed within a bin, we define the partitioning coefficient $K$ as:

$$K = \frac{N_{\text{droplet}}}{N_{\text{total}} - N_{\text{droplet}}}$$

where $N_{\text{droplet}}$ is the number of molecules within the condensate, $N_{\text{total}}$ is the total number of molecules obtained from summing time-averaged density values. Each bin number density $\rho_i$ is time-averaged over multiple time frames:

$$N_{\text{total}} = \sum_i \langle \rho_i \rangle_T$$

where $\langle \rho_i \rangle_T$ is the time-averaged density in bin $i$. The error in $N_{\text{total}}$ comes from:

$$\sigma_{\text{density}} = \sqrt{\sum_i \sigma^2_{\rho_i, T}}$$

where $\sigma_{\rho_i, T}$ is the standard deviation of density in bin $i$ over time. The error also includes the uncertainty associated with the bin slicing of the box (5%)

$$\sigma_{\text{volume}} = 0.05 \times N_{\text{total}}$$

thus

$$\sigma_{N_{\text{total}}} = \sqrt{\sigma^2_{\text{density}} + \sigma^2_{\text{volume}}}$$

The number of molecules in the detected cluster is also time-averaged:

$$N_{\text{droplet}} = \sum_j \langle \rho_j \rangle_T$$

where the sum is taken over bins belonging to the detected cluster. The error in $N_{\text{droplet}}$ accounts for density fluctuations over time

$$\sigma_{N_{\text{droplet}}} = \sqrt{\sum_j \sigma^2_{\rho_j, T}}$$

bin volume uncertainty (5% relative error):

$$\sigma_{\text{volume, cluster}} = 0.05 \times N_{\text{droplet}}$$

thus total error in $N_{\text{droplet}}$

$$\sigma_{N_{\text{droplet}}} = \sqrt{\sigma^2_{N_{\text{droplet}}, \text{density}} + \sigma^2_{\text{volume, cluster}}}.$$

Using error propagation for the fraction:

$$\sigma_K = K \times \sqrt{\left(\frac{\sigma_{N_{\text{droplet}}}}{N_{\text{droplet}}}\right)^2 + \left(\frac{\sigma_{N_{\text{total}}}}{N_{\text{total}}}\right)^2}$$

where $\sigma_K$ is the uncertainty in the partitioning coefficient.

Table S5. **Partitioning coefficient of 2-methylpropane in a system with various $(YXY)_2$ peptides computed in the all-atom simulation at 300 K.**

| Peptide | $K$ (2-methylpropane) |
|---|---|
| YFYYFY | 3.2685 ± 0.5739 |
| YNYYNY | 2.2639 ± 0.3062 |
| YYYYYY | 17.8565 ± 1.7403 |
| YAYYAY | 1.5507 ± 0.2764 |
| YLYYLY | 1.6043 ± 0.3664 |
| YWYYWY | 7.9603 ± 0.9250 |
| YSYYSY | 2.1425 ± 0.3283 |
| YCYYCY | 3.3011 ± 0.3864 |
| YQYYQY | 2.6204 ± 0.3578 |
| YGYYGY | 2.0835 ± 0.2811 |
| YHYYHY | 3.7291 ± 0.4253 |
| YIYYIY | 2.6119 ± 0.3249 |
| YMYYMY | 4.1173 ± 0.4808 |
| YPYYPY | 4.8753 ± 0.6382 |
| YTYYTY | 3.0928 ± 0.3794 |
| YVYYVY | 2.6555 ± 0.3311 |

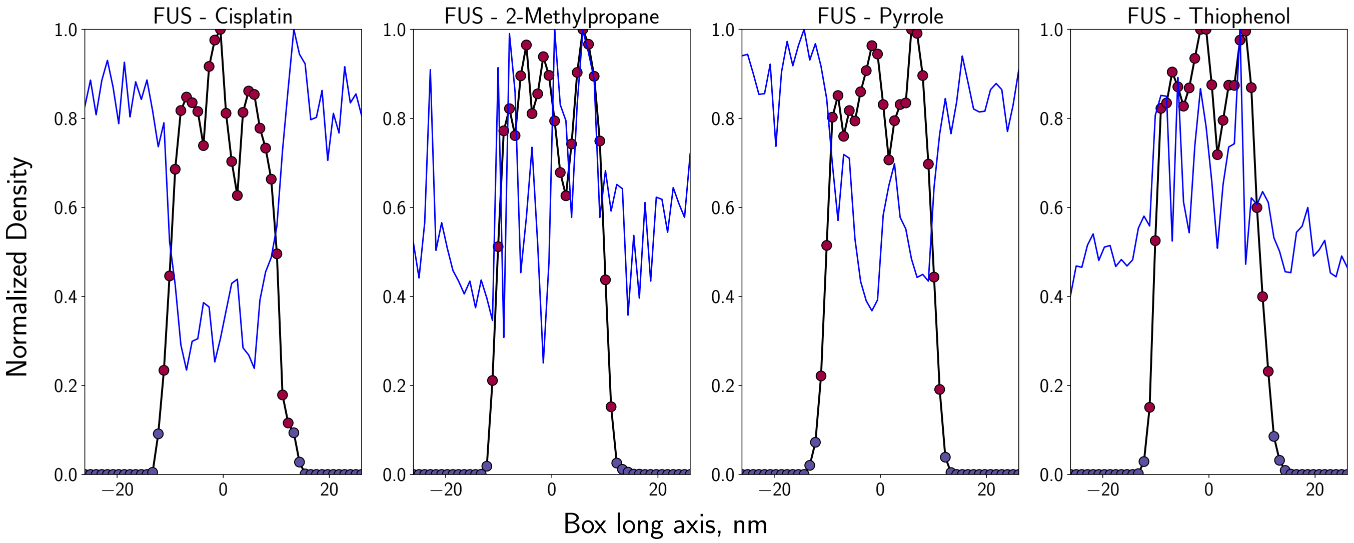


Figure S5. **Averaged density distributions of the protein (markers) and small molecules (solid blue line) for FUS LCD + small molecule systems obtained from the all-atom simulations at 300 K.** Red markers indicate the phase boundary region, within which the small molecule distribution is counted as partitioning.

#### B. Coarse-grained simulations

Table S6. System parameters for coarse-grained slab simulations. Each system is initially compressed in an $NPT$ simulation to achieve a protein density of approximately 0.3 g/cm$^3$. For peptide-based systems, the simulation box is expanded to match the molar concentration of small molecules of the corresponding all-atom system.

| Protein | Box size [Å$^3$] | $N$ beads total | $N$ SM | $N$ protein chains | $C$ SM [mM] |
|---|---|---|---|---|---|
| YMYYMY | 409.1 × 97.5 × 97.5 | 4724 | 350 | 729 | 150 |
| YGYYGY | 418.6 × 96.4 × 96.4 | 4724 | 350 | 729 | 150 |
| YTYYTY | 409.3 × 97.5 × 97.5 | 4724 | 350 | 729 | 150 |
| YAYYAY | 500.0 × 88.2 × 88.2 | 4724 | 350 | 729 | 150 |
| YYYYYY | 468.0 × 91.9 × 91.9 | 4724 | 350 | 729 | 150 |
| YVYYVY | 405.7 × 97.9 × 97.9 | 4724 | 350 | 729 | 150 |
| YLYYLY | 438.0 × 94.3 × 94.3 | 4724 | 350 | 729 | 150 |
| YQYYQY | 404.0 × 98.0 × 98.0 | 4724 | 350 | 729 | 150 |
| YWYYWY | 456.0 × 92.3 × 92.3 | 4724 | 350 | 729 | 150 |
| YFYYFY | 470.0 × 90.9 × 90.9 | 4724 | 350 | 729 | 150 |
| YSYYSY | 413.4 × 96.6 × 96.6 | 4724 | 350 | 729 | 150 |
| YHYYHY | 402.4 × 98.3 × 98.3 | 4724 | 350 | 729 | 150 |
| YNYYNY | 372.0 × 102.2 × 102.2 | 4724 | 350 | 729 | 150 |
| YPYYPY | 409.1 × 97.5 × 97.5 | 4724 | 350 | 729 | 150 |
| YCYYCY | 412.0 × 97.1 × 97.1 | 4724 | 350 | 729 | 150 |
| YIYYIY | 397.3 × 98.9 × 98.9 | 4724 | 350 | 729 | 150 |
| FWAFWF | 514.0 × 97.9 × 97.9 | 4724 | 350 | 729 | 118 |
| FYAFYF | 502.0 × 107.7 × 107.7 | 6350 | 350 | 1000 | 100 |
| NYANYN | 548.0 × 103.1 × 103.1 | 6350 | 350 | 1000 | 100 |
| NFWAFS | 572.0 × 92.7 × 92.7 | 4724 | 350 | 729 | 118 |
| YQHQHY | 523.0 × 97.0 × 97.0 | 4724 | 350 | 729 | 118 |
| FUS | 126.3 × 532.2 × 126.3 | 10910 | 150 | 64 | 30 |
| A1 | 571.5 × 106.3 × 106.3 | 8768 | 114 | 64 | 30 |
| TIA1 | 490.0 × 104.7 × 104.7 | 10910 | 95 | 64 | 30 |
| EWSR1 | 500.0 × 111.4 × 111.4 | 7560 | 110 | 27 | 30 |

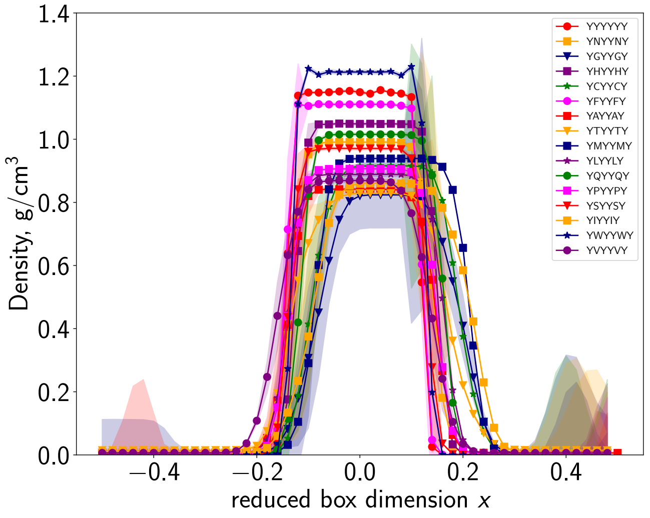


Figure S6. Density profiles of peptides (YXY)$_2$, simulated using the Mpipi model at 300 K.

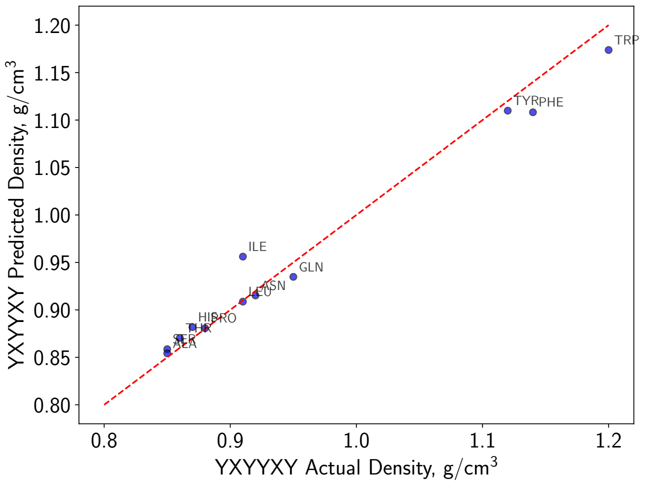


Figure S7. Predicted average density of the peptides (YXY)$_2$ with the random forest regressor; the model is used to estimate the relative density values $\rho$(YXY)$_2$/$\rho$(Y$_6$) for YKYYKY, YRYYRY, YDYYDY, and YEYYEY.

Table S7. **Density of the dense region of the $(YXY)_2$ condensates relative to the $Y_6$ condensate ($\rho_{Y_6} = 1.147$ g/cm$^3$).** The values are obtained from the coarse-grained simulation at 300 K with Mpipi force field (Fig. S6).

| Peptide | $\rho$ $(YXY)_2$ $/\rho(Y_6)$ |
|---|---|
| YMYYMY | 0.804 |
| YGYYGY | 0.703 |
| YKYYKY | 0.880 |
| YTYYTY | 0.709 |
| YRYYRY | 0.885 |
| YAYYAY | 0.723 |
| YDYYDY | 0.857 |
| YEYYEY | 0.869 |
| YYYYYY | 1.000 |
| YVYYVY | 0.747 |
| YLYYLY | 0.770 |
| YQYYQY | 0.882 |
| YWYYWY | 1.043 |
| YFYYFY | 0.967 |
| YSYYSY | 0.843 |
| YHYYHY | 0.909 |
| YNYYNY | 0.859 |
| YPYYPY | 0.781 |
| YCYYCY | 0.784 |
| YIYYIY | 0.739 |

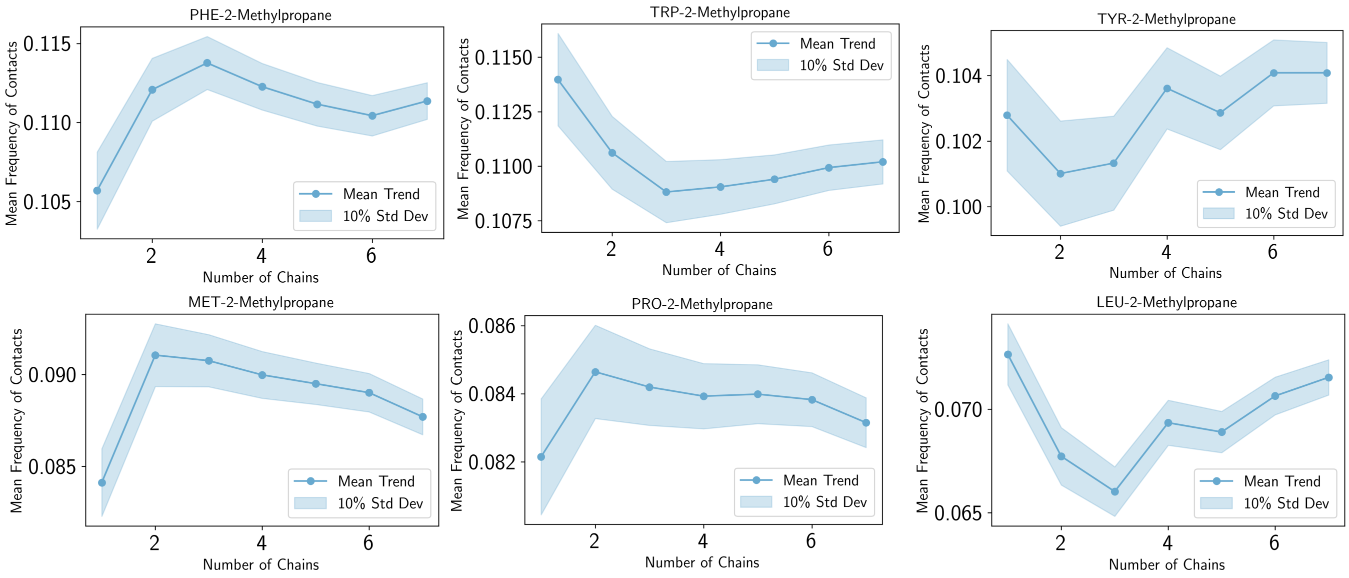


Figure S8. Cumulative average of the mean normalized frequency of contacts computed per residue, plotted as a function of the number of chains (with randomized sequences) used to compute the mean value. The cumulative average value per data point is averaged among 25 randomly permuted chains.

#### S4. MAPPS PARAMETERS FOR SMALL MOLECULES

Table S8. **Coarse-grained parameters for the non-bonded interactions between molecule–molecule pairs, modeled via the Wang–Frenkel potential**. Each molecule has a net charge $q$ of zero.

| Compound | $\varepsilon$ [kcal/mol] | $\sigma$ [Å] | $\nu$ | $\mu$ | $R_{\text{cut}}$ [Å] |
|---|---|---|---|---|---|
| 2-Methylpropane | 0.1420 | 5.3160 | 1 | 4 | 15.9480 |
| Acetic Acid | 0.1177 | 4.7340 | 1 | 3 | 14.2020 |
| Aniline | 0.0409 | 5.6540 | 1 | 4 | 16.9620 |
| Benzonitrile | 0.3457 | 5.7670 | 1 | 3 | 17.3010 |
| Cisplatin | 0.2680 | 5.6780 | 1 | 3 | 17.0340 |
| Cyclopentanol | 0.1907 | 5.6280 | 1 | 4 | 16.8840 |
| Dioxane | 0.1201 | 5.4510 | 1 | 4 | 16.3530 |
| Furan | 0.0412 | 4.9730 | 1 | 4 | 14.9190 |
| Nitrobenzene | 0.4050 | 5.8910 | 1 | 3 | 17.6730 |
| Phenol | 0.1116 | 5.5870 | 1 | 3 | 16.7610 |
| Pyrrole | 0.0709 | 5.0721 | 1 | 7 | 15.2163 |
| Styrene | 0.0769 | 5.9776 | 1 | 5 | 17.9328 |
| Thiophenol | 0.1053 | 5.7890 | 1 | 5 | 17.3670 |
| Toluene | 0.0010 | 5.7618 | 1 | 4 | 17.2854 |

Table S9: Coarse-grained parameters for the non-bonded interactions between molecule–amino acids pairs, modeled via the Wang–Frenkel potential.

| **Amino Acid** | $\varepsilon$ [kcal/mol] | $\sigma$ [Å] | $\nu$ | $\mu$ | $R_{\text{cut}}$ [Å] |
|---|---|---|---|---|---|
| **2-Methylpropane** | | | | | |
| MET | 0.6714 | 5.8920 | 1 | 3 | 17.6759 |
| GLY | 0.3654 | 5.0056 | 1 | 3 | 15.0167 |
| LYS | 0.4044 | 5.9937 | 1 | 3 | 17.9810 |
| THR | 0.3706 | 5.6025 | 1 | 3 | 16.8076 |
| ARG | 0.6005 | 6.0775 | 1 | 3 | 18.2326 |
| ALA | 0.2988 | 5.2930 | 1 | 3 | 15.8791 |
| ASP | 0.2824 | 5.5698 | 1 | 3 | 16.7093 |

| Continued from previous page | | | | | |
|---|---|---|---|---|---|
| **Amino Acid** | $\varepsilon$ | $\sigma$ | $\nu$ | $\mu$ | $R_{cut}$ |
| GLU | 0.3382 | 5.7468 | 1 | 3 | 17.2405 |
| TYR | 0.9911 | 6.0248 | 1 | 3 | 18.0745 |
| VAL | 0.4763 | 5.7910 | 1 | 3 | 17.3730 |
| LEU | 0.5245 | 5.9250 | 1 | 3 | 17.7751 |
| GLN | 0.3238 | 5.7969 | 1 | 3 | 17.3908 |
| TRP | 1.0945 | 6.1913 | 1 | 3 | 18.5738 |
| PHE | 1.0255 | 5.9728 | 1 | 3 | 17.9183 |
| SER | 0.3725 | 5.3643 | 1 | 3 | 16.0930 |
| HIS | 0.5770 | 5.8269 | 1 | 3 | 17.4807 |
| ASN | 0.4487 | 5.6197 | 1 | 3 | 16.8590 |
| PRO | 0.6184 | 5.5611 | 1 | 3 | 16.6832 |
| CYS | 0.3767 | 5.5202 | 1 | 3 | 16.5605 |
| ILE | 0.4855 | 6.1188 | 1 | 3 | 18.3565 |
| **Acetic Acid** | | | | | |
| MET | 0.1995 | 5.6010 | 1 | 3 | 16.8029 |
| GLY | 0.1224 | 4.7146 | 1 | 3 | 14.1437 |
| LYS | 0.1827 | 5.7027 | 1 | 3 | 17.1080 |
| THR | 0.1595 | 5.3115 | 1 | 3 | 15.9346 |
| ARG | 0.2022 | 5.7865 | 1 | 3 | 17.3596 |
| ALA | 0.1263 | 5.0020 | 1 | 3 | 15.0061 |
| ASP | 0.1549 | 5.2788 | 1 | 3 | 15.8363 |
| GLU | 0.1676 | 5.4558 | 1 | 3 | 16.3675 |
| TYR | 0.3333 | 5.7338 | 1 | 3 | 17.2015 |
| VAL | 0.1332 | 5.5000 | 1 | 3 | 16.5000 |
| LEU | 0.1906 | 5.6340 | 1 | 3 | 16.9021 |
| GLN | 0.2165 | 5.5059 | 1 | 3 | 16.5178 |
| TRP | 0.2668 | 5.9003 | 1 | 3 | 17.7008 |
| PHE | 0.2398 | 5.6818 | 1 | 3 | 17.0453 |
| SER | 0.1416 | 5.0733 | 1 | 3 | 15.2200 |
| HIS | 0.2314 | 5.5359 | 1 | 3 | 16.6077 |
| ASN | 0.1563 | 5.3287 | 1 | 3 | 15.9860 |
| PRO | 0.1228 | 5.2701 | 1 | 3 | 15.8102 |
| CYS | 0.1345 | 5.2292 | 1 | 3 | 15.6875 |
| ILE | 0.1585 | 5.8278 | 1 | 3 | 17.4835 |
| **Aniline** | | | | | |
| MET | 0.5910 | 6.0610 | 1 | 3 | 18.1829 |
| GLY | 0.3352 | 5.1746 | 1 | 3 | 15.5237 |
| LYS | 0.4926 | 6.1627 | 1 | 3 | 18.4880 |
| THR | 0.3090 | 5.7715 | 1 | 3 | 17.3146 |
| ARG | 0.6720 | 6.2465 | 1 | 3 | 18.7396 |
| ALA | 0.3721 | 5.4620 | 1 | 3 | 16.3861 |
| ASP | 0.4090 | 5.7388 | 1 | 3 | 17.2163 |
| GLU | 0.3939 | 5.9158 | 1 | 3 | 17.7475 |
| TYR | 0.8000 | 6.1938 | 1 | 3 | 18.5815 |
| VAL | 0.3512 | 5.9600 | 1 | 3 | 17.8800 |
| LEU | 0.4354 | 6.0940 | 1 | 3 | 18.2821 |
| GLN | 0.4518 | 5.9659 | 1 | 3 | 17.8978 |
| TRP | 0.8421 | 6.3603 | 1 | 3 | 19.0808 |
| PHE | 0.5912 | 6.1418 | 1 | 3 | 18.4253 |
| SER | 0.4646 | 5.5333 | 1 | 3 | 16.6000 |
| HIS | 0.5676 | 5.9959 | 1 | 3 | 17.9877 |
| ASN | 0.5398 | 5.7887 | 1 | 3 | 17.3660 |
| PRO | 0.4250 | 5.7301 | 1 | 3 | 17.1902 |
| CYS | 0.3604 | 5.6892 | 1 | 3 | 17.0675 |
| ILE | 0.4222 | 6.2878 | 1 | 3 | 18.8635 |
| **Benzonitrile** | | | | | |
| MET | 0.5724 | 6.1170 | 1 | 3 | 18.3509 |
| GLY | 0.3659 | 5.2306 | 1 | 3 | 15.6917 |
| LYS | 0.4383 | 6.2187 | 1 | 3 | 18.6560 |

Continued from previous page

| **Amino Acid** | $\varepsilon$ | $\sigma$ | $\nu$ | $\mu$ | $R_{cut}$ |
|---|---|---|---|---|---|
| THR | 0.3730 | 5.8275 | 1 | 3 | 17.4826 |
| ARG | 0.5891 | 6.3025 | 1 | 3 | 18.9076 |
| ALA | 0.3026 | 5.5180 | 1 | 3 | 16.5541 |
| ASP | 0.3061 | 5.7948 | 1 | 3 | 17.3843 |
| GLU | 0.3286 | 5.9718 | 1 | 3 | 17.9155 |
| TYR | 0.9000 | 6.2498 | 1 | 3 | 18.7495 |
| VAL | 0.4883 | 6.0160 | 1 | 3 | 18.0480 |
| LEU | 0.5429 | 6.1500 | 1 | 3 | 18.4501 |
| GLN | 0.5292 | 6.0219 | 1 | 3 | 18.0658 |
| TRP | 0.9265 | 6.4163 | 1 | 3 | 19.2488 |
| PHE | 0.9702 | 6.1978 | 1 | 3 | 18.5933 |
| SER | 0.4034 | 5.5893 | 1 | 3 | 16.7680 |
| HIS | 0.7072 | 6.0519 | 1 | 3 | 18.1557 |
| ASN | 0.3836 | 5.8447 | 1 | 3 | 17.5340 |
| PRO | 0.4640 | 5.7861 | 1 | 3 | 17.3582 |
| CYS | 0.3010 | 5.7452 | 1 | 3 | 17.2355 |
| ILE | 0.4293 | 6.3438 | 1 | 3 | 19.0315 |
| **Cisplatin** | | | | | |
| MET | 0.3333 | 6.0730 | 1 | 3 | 18.2189 |
| GLY | 0.2804 | 5.1866 | 1 | 3 | 15.5597 |
| LYS | 0.4679 | 6.1747 | 1 | 3 | 18.5240 |
| THR | 0.2609 | 5.7835 | 1 | 3 | 17.3506 |
| ARG | 0.4885 | 6.2585 | 1 | 3 | 18.7756 |
| ALA | 0.1700 | 5.4740 | 1 | 3 | 16.4221 |
| ASP | 0.1991 | 5.7508 | 1 | 3 | 17.2523 |
| GLU | 0.2358 | 5.9278 | 1 | 3 | 17.7835 |
| TYR | 0.5000 | 6.2058 | 1 | 3 | 18.6175 |
| VAL | 0.2394 | 5.9720 | 1 | 3 | 17.9160 |
| LEU | 0.2630 | 6.1060 | 1 | 3 | 18.3181 |
| GLN | 0.3501 | 5.9779 | 1 | 3 | 17.9338 |
| TRP | 0.6413 | 6.3723 | 1 | 3 | 19.1168 |
| PHE | 0.4727 | 6.1538 | 1 | 3 | 18.4613 |
| SER | 0.2393 | 5.5453 | 1 | 3 | 16.6360 |
| HIS | 0.4317 | 6.0079 | 1 | 3 | 18.0237 |
| ASN | 0.3400 | 5.8007 | 1 | 3 | 17.4020 |
| PRO | 0.2291 | 5.7421 | 1 | 3 | 17.2262 |
| CYS | 0.2094 | 5.7012 | 1 | 3 | 17.1035 |
| ILE | 0.3163 | 6.2998 | 1 | 3 | 18.8995 |
| **Cyclopentanol** | | | | | |
| MET | 0.5981 | 6.0480 | 1 | 3 | 18.1439 |
| GLY | 0.2964 | 5.1616 | 1 | 3 | 15.4847 |
| LYS | 0.5092 | 6.1497 | 1 | 3 | 18.4490 |
| THR | 0.3696 | 5.7585 | 1 | 3 | 17.2756 |
| ARG | 0.7600 | 6.2335 | 1 | 3 | 18.7006 |
| ALA | 0.3073 | 5.4490 | 1 | 3 | 16.3471 |
| ASP | 0.3162 | 5.7258 | 1 | 3 | 17.1773 |
| GLU | 0.4428 | 5.9028 | 1 | 3 | 17.7085 |
| TYR | 0.8000 | 6.1808 | 1 | 3 | 18.5425 |
| VAL | 0.4602 | 5.9470 | 1 | 3 | 17.8410 |
| LEU | 0.4482 | 6.0810 | 1 | 3 | 18.2431 |
| GLN | 0.5076 | 5.9529 | 1 | 3 | 17.8588 |
| TRP | 1.0521 | 6.3473 | 1 | 3 | 19.0418 |
| PHE | 0.9059 | 6.1288 | 1 | 3 | 18.3863 |
| SER | 0.3487 | 5.5203 | 1 | 3 | 16.5610 |
| HIS | 0.6549 | 5.9829 | 1 | 3 | 17.9487 |
| ASN | 0.4880 | 5.7757 | 1 | 3 | 17.3270 |
| PRO | 0.4717 | 5.7171 | 1 | 3 | 17.1512 |
| CYS | 0.3637 | 5.6762 | 1 | 3 | 17.0285 |
| ILE | 0.5318 | 6.2748 | 1 | 3 | 18.8245 |

Continued from previous page

| Amino Acid | $\varepsilon$ | $\sigma$ | $\nu$ | $\mu$ | $R_{\text{cut}}$ |
|---|---|---|---|---|---|
| **Dioxane** | | | | | |
| MET | 0.5527 | 5.9595 | 1 | 3 | 17.8784 |
| GLY | 0.2885 | 5.0731 | 1 | 3 | 15.2192 |
| LYS | 0.4166 | 6.0612 | 1 | 3 | 18.1835 |
| THR | 0.3249 | 5.6700 | 1 | 3 | 17.0101 |
| ARG | 0.5808 | 6.1450 | 1 | 3 | 18.4351 |
| ALA | 0.3380 | 5.3605 | 1 | 3 | 16.0816 |
| ASP | 0.3282 | 5.6373 | 1 | 3 | 16.9118 |
| GLU | 0.3008 | 5.8143 | 1 | 3 | 17.4430 |
| TYR | 0.7333 | 6.0923 | 1 | 3 | 18.2770 |
| VAL | 0.4629 | 5.8585 | 1 | 3 | 17.5755 |
| LEU | 0.6089 | 5.9925 | 1 | 3 | 17.9776 |
| GLN | 0.2399 | 5.8644 | 1 | 3 | 17.5933 |
| TRP | 1.0381 | 6.2588 | 1 | 3 | 18.7763 |
| PHE | 0.9334 | 6.0403 | 1 | 3 | 18.1208 |
| SER | 0.3443 | 5.4318 | 1 | 3 | 16.2955 |
| HIS | 0.5416 | 5.8944 | 1 | 3 | 17.6832 |
| ASN | 0.5127 | 5.6872 | 1 | 3 | 17.0615 |
| PRO | 0.3963 | 5.6286 | 1 | 3 | 16.8857 |
| CYS | 0.2920 | 5.5877 | 1 | 3 | 16.7630 |
| ILE | 0.3300 | 6.1863 | 1 | 3 | 18.5590 |
| **Furan** | | | | | |
| MET | 0.3992 | 5.7205 | 1 | 3 | 17.1614 |
| GLY | 0.2827 | 4.8341 | 1 | 3 | 14.5022 |
| LYS | 0.3800 | 5.8222 | 1 | 3 | 17.4665 |
| THR | 0.3429 | 5.4310 | 1 | 3 | 16.2931 |
| ARG | 0.4514 | 5.9060 | 1 | 3 | 17.7181 |
| ALA | 0.3158 | 5.1215 | 1 | 3 | 15.3646 |
| ASP | 0.3126 | 5.3983 | 1 | 3 | 16.1948 |
| GLU | 0.3232 | 5.5753 | 1 | 3 | 16.7260 |
| TYR | 0.8042 | 5.8533 | 1 | 3 | 17.5600 |
| VAL | 0.3566 | 5.6195 | 1 | 3 | 16.8585 |
| LEU | 0.4998 | 5.7535 | 1 | 3 | 17.2606 |
| GLN | 0.4902 | 5.6254 | 1 | 3 | 16.8763 |
| TRP | 0.9893 | 6.0198 | 1 | 3 | 18.0593 |
| PHE | 0.5600 | 5.8013 | 1 | 3 | 17.4038 |
| SER | 0.3710 | 5.1928 | 1 | 3 | 15.5785 |
| HIS | 0.6524 | 5.6554 | 1 | 3 | 16.9662 |
| ASN | 0.4145 | 5.4482 | 1 | 3 | 16.3445 |
| PRO | 0.4264 | 5.3896 | 1 | 3 | 16.1687 |
| CYS | 0.2988 | 5.3487 | 1 | 3 | 16.0460 |
| ILE | 0.3721 | 5.9473 | 1 | 3 | 17.8420 |
| **Nitrobenzene** | | | | | |
| MET | 0.5912 | 6.1794 | 1 | 3 | 18.5383 |
| GLY | 0.3415 | 5.2930 | 1 | 3 | 15.8790 |
| LYS | 0.4398 | 6.2811 | 1 | 3 | 18.8434 |
| THR | 0.3016 | 5.8900 | 1 | 3 | 17.6699 |
| ARG | 0.5908 | 6.3650 | 1 | 3 | 19.0949 |
| ALA | 0.2992 | 5.5805 | 1 | 3 | 16.7415 |
| ASP | 0.3452 | 5.8572 | 1 | 3 | 17.5716 |
| GLU | 0.3752 | 6.0343 | 1 | 3 | 18.1029 |
| TYR | 0.9737 | 6.3123 | 1 | 3 | 18.9368 |
| VAL | 0.4135 | 6.0784 | 1 | 3 | 18.2353 |
| LEU | 0.4019 | 6.2125 | 1 | 3 | 18.6375 |
| GLN | 0.4854 | 6.0844 | 1 | 3 | 18.2531 |
| TRP | 1.1112 | 6.4787 | 1 | 3 | 19.4362 |
| PHE | 0.8515 | 6.2602 | 1 | 3 | 18.7807 |
| SER | 0.5117 | 5.6518 | 1 | 3 | 16.9554 |
| HIS | 0.6385 | 6.1143 | 1 | 3 | 18.3430 |

Continued from previous page

| Amino Acid | $\varepsilon$ | $\sigma$ | $\nu$ | $\mu$ | $R_{cut}$ |
|---|---|---|---|---|---|
| ASN | 0.4402 | 5.9071 | 1 | 3 | 17.7214 |
| PRO | 0.4496 | 5.8485 | 1 | 3 | 17.5456 |
| CYS | 0.4039 | 5.8076 | 1 | 3 | 17.4229 |
| ILE | 0.2932 | 6.4063 | 1 | 3 | 19.2189 |
| **Phenol** | | | | | |
| MET | 0.5832 | 6.0275 | 1 | 3 | 18.0824 |
| GLY | 0.2944 | 5.1411 | 1 | 3 | 15.4232 |
| LYS | 0.5094 | 6.1292 | 1 | 3 | 18.3875 |
| THR | 0.3854 | 5.7380 | 1 | 3 | 17.2141 |
| ARG | 0.6689 | 6.2130 | 1 | 3 | 18.6391 |
| ALA | 0.2861 | 5.4285 | 1 | 3 | 16.2856 |
| ASP | 0.2994 | 5.7053 | 1 | 3 | 17.1158 |
| GLU | 0.3900 | 5.8823 | 1 | 3 | 17.6470 |
| TYR | 0.8438 | 6.1603 | 1 | 3 | 18.4810 |
| VAL | 0.4853 | 5.9265 | 1 | 3 | 17.7795 |
| LEU | 0.4965 | 6.0605 | 1 | 3 | 18.1816 |
| GLN | 0.4830 | 5.9324 | 1 | 3 | 17.7973 |
| TRP | 0.9427 | 6.3268 | 1 | 3 | 18.9803 |
| PHE | 0.8491 | 6.1083 | 1 | 3 | 18.3248 |
| SER | 0.4357 | 5.4998 | 1 | 3 | 16.4995 |
| HIS | 0.6837 | 5.9624 | 1 | 3 | 17.8872 |
| ASN | 0.4391 | 5.7552 | 1 | 3 | 17.2655 |
| PRO | 0.6313 | 5.6966 | 1 | 3 | 17.0897 |
| CYS | 0.4520 | 5.6557 | 1 | 3 | 16.9670 |
| ILE | 0.4560 | 6.2543 | 1 | 3 | 18.7630 |
| **Pyrrole** | | | | | |
| MET | 0.4015 | 5.7700 | 1 | 3 | 17.3101 |
| GLY | 0.1530 | 4.8836 | 1 | 3 | 14.6508 |
| LYS | 0.2991 | 5.8717 | 1 | 3 | 17.6152 |
| THR | 0.2204 | 5.4806 | 1 | 3 | 16.4417 |
| ARG | 0.4973 | 5.9556 | 1 | 3 | 17.8667 |
| ALA | 0.1807 | 5.1711 | 1 | 3 | 15.5133 |
| ASP | 0.2580 | 5.4478 | 1 | 3 | 16.3434 |
| GLU | 0.3346 | 5.6249 | 1 | 3 | 16.8747 |
| TYR | 0.6333 | 5.9029 | 1 | 3 | 17.7086 |
| VAL | 0.3163 | 5.6690 | 1 | 3 | 17.0071 |
| LEU | 0.2562 | 5.8031 | 1 | 3 | 17.4093 |
| GLN | 0.3610 | 5.6750 | 1 | 3 | 17.0249 |
| TRP | 0.7736 | 6.0693 | 1 | 3 | 18.2080 |
| PHE | 0.3927 | 5.8508 | 1 | 3 | 17.5525 |
| SER | 0.3210 | 5.2424 | 1 | 3 | 15.7272 |
| HIS | 0.4732 | 5.7049 | 1 | 3 | 17.1148 |
| ASN | 0.3477 | 5.4977 | 1 | 3 | 16.4932 |
| PRO | 0.3110 | 5.4391 | 1 | 3 | 16.3174 |
| CYS | 0.2844 | 5.3982 | 1 | 3 | 16.1947 |
| ILE | 0.3265 | 5.9969 | 1 | 3 | 17.9907 |
| **Styrene** | | | | | |
| MET | 0.7437 | 6.2228 | 1 | 3 | 18.6683 |
| GLY | 0.3561 | 5.3364 | 1 | 3 | 16.0091 |
| LYS | 0.5460 | 6.3245 | 1 | 3 | 18.9734 |
| THR | 0.3551 | 5.9333 | 1 | 3 | 17.8000 |
| ARG | 0.8885 | 6.4083 | 1 | 3 | 19.2250 |
| ALA | 0.5143 | 5.6238 | 1 | 3 | 16.8715 |
| ASP | 0.2951 | 5.9006 | 1 | 3 | 17.7017 |
| GLU | 0.3981 | 6.0776 | 1 | 3 | 18.2329 |
| TYR | 0.9368 | 6.3556 | 1 | 3 | 19.0669 |
| VAL | 0.6368 | 6.1218 | 1 | 3 | 18.3654 |
| LEU | 0.8555 | 6.2558 | 1 | 3 | 18.7675 |
| GLN | 0.7011 | 6.1277 | 1 | 3 | 18.3832 |

Continued from previous page

| Amino Acid | $\varepsilon$ | $\sigma$ | $\nu$ | $\mu$ | $R_{\text{cut}}$ |
|---|---|---|---|---|---|
| TRP | 1.1470 | 6.5221 | 1 | 3 | 19.5662 |
| PHE | 0.8861 | 6.3036 | 1 | 3 | 18.9107 |
| SER | 0.6475 | 5.6951 | 1 | 3 | 17.0854 |
| HIS | 0.9154 | 6.1577 | 1 | 3 | 18.4731 |
| ASN | 0.5515 | 5.9505 | 1 | 3 | 17.8514 |
| PRO | 0.7029 | 5.8919 | 1 | 3 | 17.6756 |
| CYS | 0.4907 | 5.8510 | 1 | 3 | 17.5529 |
| ILE | 0.6476 | 6.4496 | 1 | 3 | 19.3489 |
| **Thiophenol** | | | | | |
| MET | 0.5650 | 6.1285 | 1 | 3 | 18.3854 |
| GLY | 0.3140 | 5.2421 | 1 | 3 | 15.7262 |
| LYS | 0.5139 | 6.2302 | 1 | 3 | 18.6905 |
| THR | 0.3941 | 5.8390 | 1 | 3 | 17.5171 |
| ARG | 0.6882 | 6.3140 | 1 | 3 | 18.9421 |
| ALA | 0.3958 | 5.5295 | 1 | 3 | 16.5886 |
| ASP | 0.2890 | 5.8063 | 1 | 3 | 17.4188 |
| GLU | 0.4259 | 5.9833 | 1 | 3 | 17.9500 |
| TYR | 0.9333 | 6.2613 | 1 | 3 | 18.7840 |
| VAL | 0.6037 | 6.0275 | 1 | 3 | 18.0825 |
| LEU | 0.5306 | 6.1615 | 1 | 3 | 18.4846 |
| GLN | 0.5980 | 6.0334 | 1 | 3 | 18.1003 |
| TRP | 1.2693 | 6.4278 | 1 | 3 | 19.2833 |
| PHE | 0.8561 | 6.2093 | 1 | 3 | 18.6278 |
| SER | 0.5122 | 5.6008 | 1 | 3 | 16.8025 |
| HIS | 0.7417 | 6.0634 | 1 | 3 | 18.1902 |
| ASN | 0.5211 | 5.8562 | 1 | 3 | 17.5685 |
| PRO | 0.6928 | 5.7976 | 1 | 3 | 17.3927 |
| CYS | 0.5961 | 5.7567 | 1 | 3 | 17.2700 |
| ILE | 0.4291 | 6.3553 | 1 | 3 | 19.0660 |
| **Toluene** | | | | | |
| MET | 0.5715 | 6.1149 | 1 | 3 | 18.3446 |
| GLY | 0.2975 | 5.2285 | 1 | 3 | 15.6854 |
| LYS | 0.2656 | 6.2166 | 1 | 3 | 18.6497 |
| THR | 0.2246 | 5.8254 | 1 | 3 | 17.4763 |
| ARG | 0.4554 | 6.3004 | 1 | 3 | 18.9013 |
| ALA | 0.3391 | 5.5159 | 1 | 3 | 16.5478 |
| ASP | 0.3144 | 5.7927 | 1 | 3 | 17.3780 |
| GLU | 0.4096 | 5.9697 | 1 | 3 | 17.9092 |
| TYR | 0.9000 | 6.2477 | 1 | 3 | 18.7432 |
| VAL | 0.5984 | 6.0139 | 1 | 3 | 18.0417 |
| LEU | 0.5393 | 6.1479 | 1 | 3 | 18.4438 |
| GLN | 0.4555 | 6.0198 | 1 | 3 | 18.0595 |
| TRP | 0.9871 | 6.4142 | 1 | 3 | 19.2425 |
| PHE | 0.7492 | 6.1957 | 1 | 3 | 18.5870 |
| SER | 0.4373 | 5.5872 | 1 | 3 | 16.7617 |
| HIS | 0.5922 | 6.0498 | 1 | 3 | 18.1494 |
| ASN | 0.4973 | 5.8426 | 1 | 3 | 17.5277 |
| PRO | 0.5730 | 5.7840 | 1 | 3 | 17.3519 |
| CYS | 0.3901 | 5.7431 | 1 | 3 | 17.2292 |
| ILE | 0.5819 | 6.3417 | 1 | 3 | 19.0252 |

#### S5. PEPTIDES + SMALL MOLECULES: ADDITIONAL RESULTS

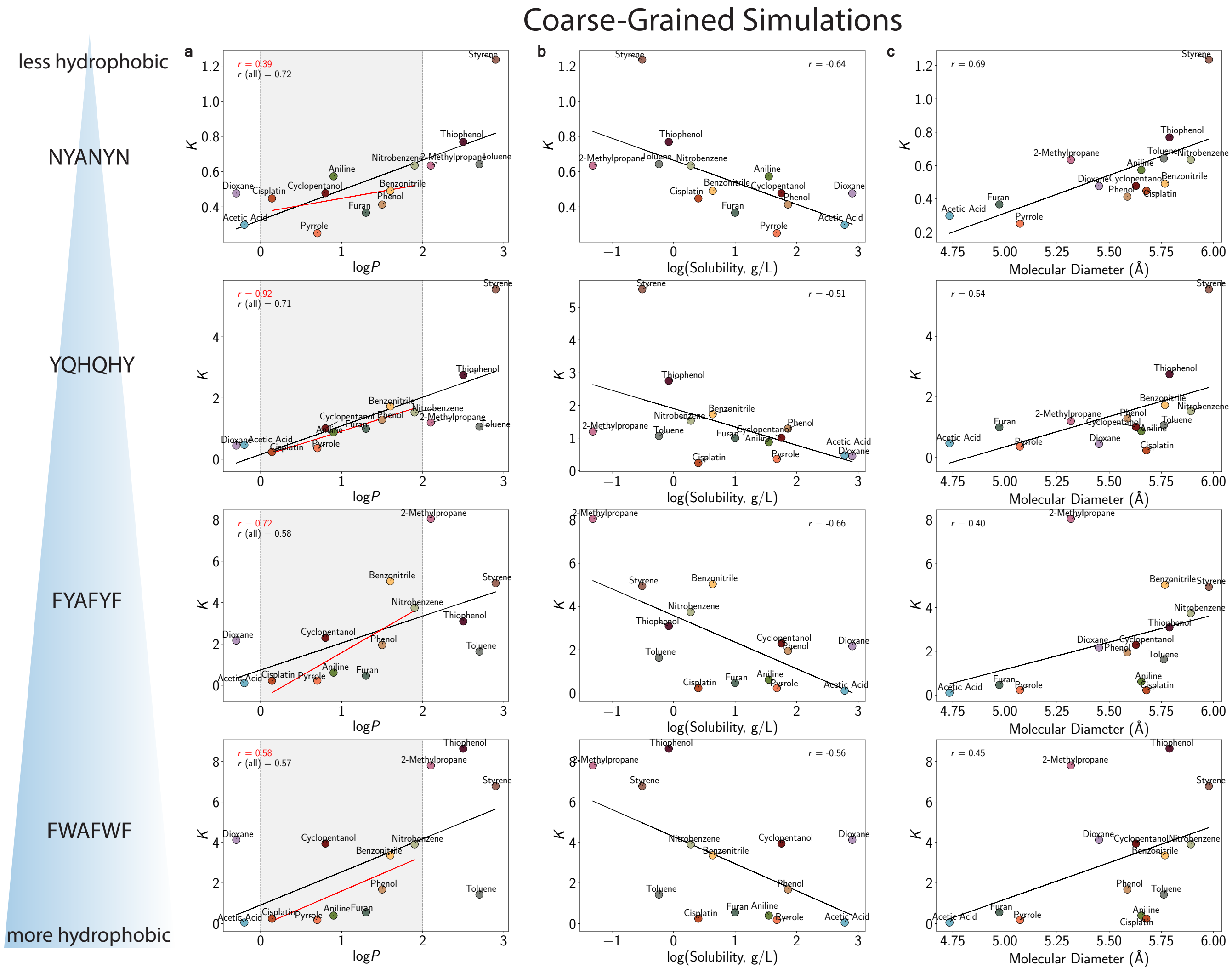


Figure S9. **Coarse-grained simulations: partitioning coefficient of a set of small molecules in NYANYN, YQHQHY, FYAFYF, and FWAFWF peptide-based condensates at 300 K**. The values are plotted against the compounds log$P$ values (a), solubility ($\log_{10}$) at 25°C [3] (b), and molecular diameter (c). The solid black line shows a linear fit with a corresponding Pearson correlation coefficient, and the red solid line shows a linear fit in the region of $0 < \log P < 2$.

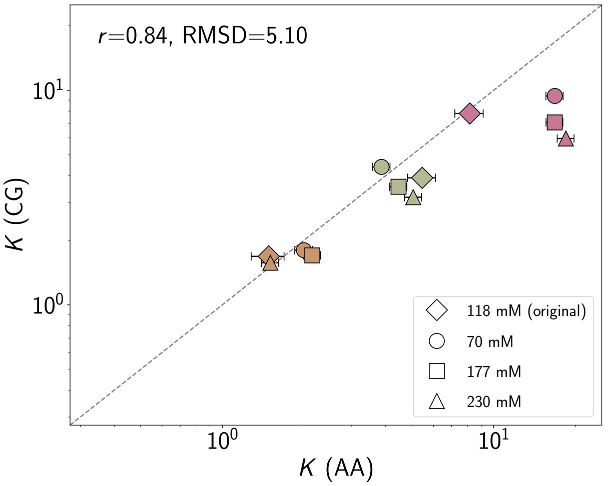


Figure S10. **Partitioning coefficients at 300 K of a subset of small molecules (2-methylpropane [pink], phenol [brown], and nitrobenzene [sage]) in the coarse-grained simulation versus all-atom simulation based on the parameterized models, for the system of FWAFWF peptides, computed for different concentrations of the small molecules (displayed in the legend).** The Pearson correlation coefficient $r$ and the root-mean-square deviation (RMSD) values are displayed. The dashed line represents the equity line.

#### S6. LCDS + SMALL MOLECULES: ADDITIONAL RESULTS

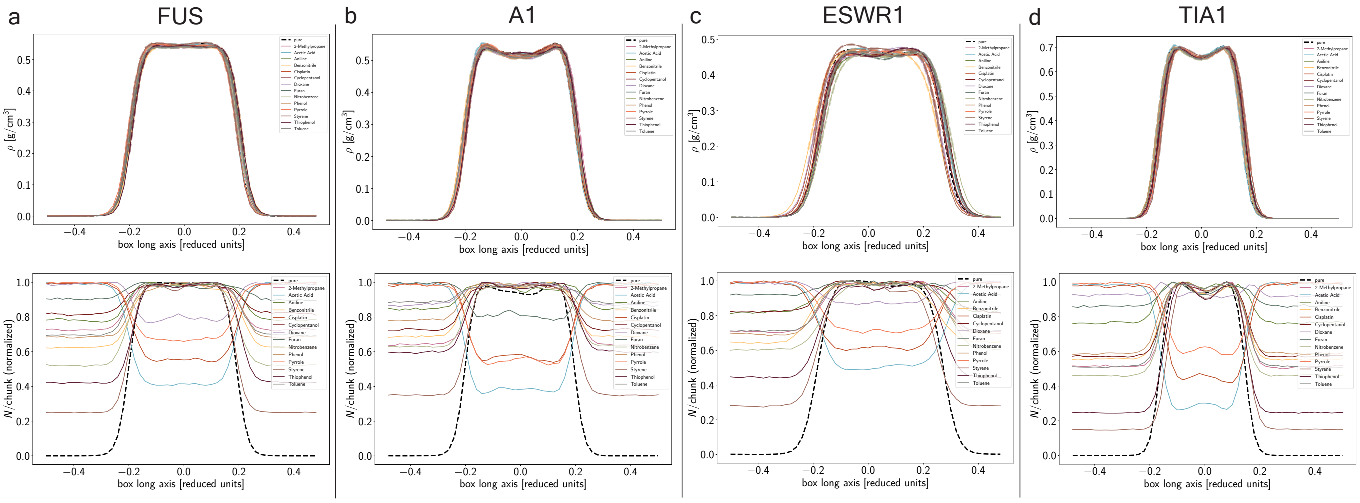


Figure S11. **Coarse-grained simulations of LCDs and small molecules at 300 K**. Density distribution of FUS (a), A1 (b), ESWR1 (c), TIA1 (d) LCDs condensates (top panel) and the relative number density (normalized by maximum value) of the pure condensate system and the protein + small molecules (bottom panel) in coarse-grained simulation at 300 K.

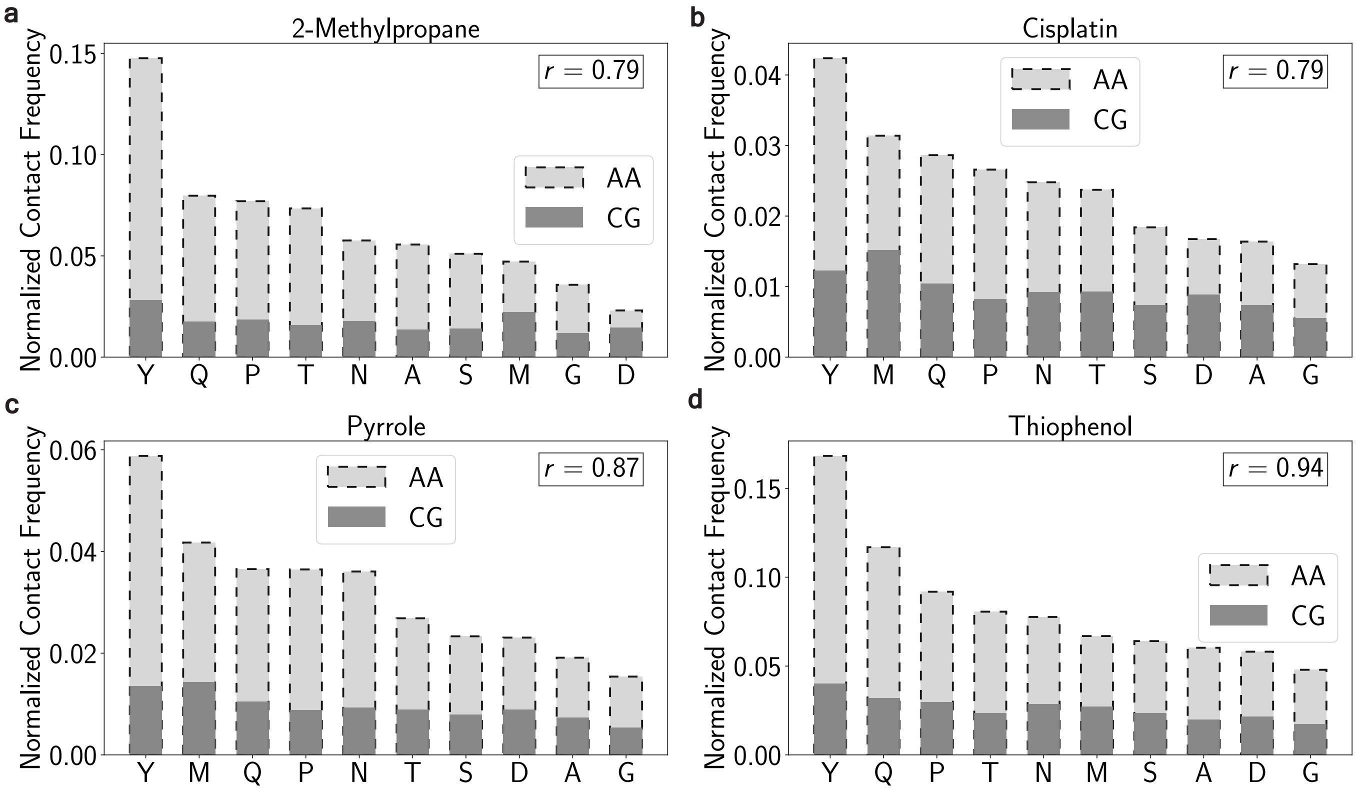


Figure S12. **Distribution of the number of contacts between the small molecules and FUS LCD condensate in the all-atom and coarse-grained simulations at 300 K.**


[1] D. Van Der Spoel, E. Lindahl, B. Hess, G. Groenhof, A. E. Mark, and H. J. Berendsen, GROMACS: fast, flexible, and free, J. Comput. Chem **26**, 1701 (2005).

[2] R. B. Best, W. Zheng, and J. Mittal, Balanced protein–water interactions improve properties of disordered proteins and non-specific protein association, J. Chem. Theory Comput. **10**, 5113 (2014).

[3] S. H. Yalkowsky, Y. He, and P. Jain, *Handbook of aqueous solubility data* (CRC press, 2016).
